# Robust and interpretable prediction of gene markers and cell types from spatial transcriptomics data

**DOI:** 10.1101/2023.05.14.540710

**Authors:** Xiao Tan, Onkar Mulay, Jacky Xie, Samual MacDonald, Taehyun Kim, Nan Ye, Peter T Simpson, Fred Roosta, Maciej Trzaskowski, Quan Nguyen

## Abstract

Spatial transcriptomic (ST) imaging and sequencing data enable us to link tissue morphological features with thousands of previously unseen gene expression values, opening a new horizon for understanding tissue biology and achieving breakthroughs in digital pathology. Deep learning models are emerging to predict gene expression or classify cell types using images as the sole input. Such models hold significant potential for clinical applications, but require improvements in interpretability and robustness. We developed STimage as a comprehensive suite of models for both regression (predicting gene expression) and classification (mapping tissue regions and cell types) tasks. STimage is the first to thoroughly address robustness (uncertainty) and interpretability. For robustness, STimage predicts gene expression based on parameter distributions rather than fixed data points, allowing for generalisation at a population scale. STimage estimates uncertainty from the data (aleatoric) and from the model (epistemic) for each of thousands of imaging tiles. STimage achieves interpretability by analysing model attribution at a single-cell level, and in the context of histopathological annotation. While existing models focus on predicting highly variable genes, STimage predicts functional genes and identifies highly predictable genes. Using diverse datasets from three cancers and one chronic disease, we assessed the model’s performance on in-distribution and out-of-distribution samples. STimage is robust to technical variations across platforms, data types, sample preservation methods, and disease types. Further, we implemented an ensemble approach, incorporating pre-trained foundation models, to improve performance and reliability, especially in cases with small training datasets. With single-cell resolution Xenium data, STimage could classify cell types for millions of individual cells. Applying STimage to proteomics data such as CODEX, we found that STimage can predict gene expression consistent with protein expression patterns. Finally, we showed that using STimage-predicted values based solely on imaging input, we could stratify patient survival groups. Overall, STimage advances spatial transcriptomics by improving the prediction of gene expression from traditional histopathological images, making it more accessible for tissue biology research and digital pathology applications.

## 1 Introduction

Hematoxylin and Eosin (H&E) staining of tissue samples has been used by pathologists for more than a century. However, microscopic examination of H&E images is often time-consuming and variable^1^. Moreover, a range of molecularly distinct cell types may not be distinguishable by eye^2^. Recent FDA approvals^3^ of whole-slide imaging (WSI) for primary diagnostic use have encouraged more widespread adoption of digital WSI^4^. However, the development and optimisation of automated cancer classification models requires a large number of pathologist-annotated tissue images^5,6^. HE2RNA^7^ was among the pioneers to train a model without pathological annotation. Instead, HE2RNA uses matched H&E images and RNA-seq data from the TCGA database for 8,725 patients across 28 different cancer types, to predict gene expression at the tile level (each WSI H&E image is split into multiple tiles). However, due to the nature of the data input, HE2RNA lacks ground truth at the tile level, using instead the aggregated gene expression value from the whole image (i.e., all tiles in the image have the same ground truth for one gene), thus limiting its ability to assess tumour heterogeneity.

Spatial transcriptomics (ST-seq) produces both imaging and sequencing information and has the unique capability of measuring over 20,000 genes without tissue dissociation, preserving tissue anatomy and the microenvironmental context^8^. However, spatial transcriptomics remains prohibitively expensive for clinical applications. To improve scalability, researchers have attempted to predict spatial gene expression using deep learning methods, including STnet^9^, Hist2Gene^10^, Hist2ST^11^, and DeepSpace^12^. However, these methods are still limited in interpretability, robustness, and accuracy. Crucially, clinical applications require models to be understandable to clinicians and patients for safety, trust, validation, and therapeutic decision-making^13^.

Here, we report STimage, a deep learning probabilistic framework with reduced uncertainty and improved robustness and interpretability, applicable even in cases where training datasets are limited. We tested STimage across three different cancer datasets, including breast cancer, skin cancer, and kidney cancer, as well as a liver immune disease dataset, assessing the performance of predicting the most variable genes or functional genes. The shared biological relationships between these genes are learnt simultaneously, allowing us to assess the performance of gene panels. We introduced a geometric assessment metric that better represents the model’s ability to predict spatial patterns.

## 2 Materials and methods

### 2.1 Datasets

The training step requires spatial transcriptomics data as input, with large histology images cropped into 299 × 299 pixel tiles. Spatial coordinates encoded by spatial barcodes for each spot, and gene expression values for each matched spot, are used as labels to train the model. The number of microns corresponding to 299 pixels depends on the image resolution and the size of the Visium spots. The tiling process ensures that each tile contains one spot of 55 µm for 10x Visium and 100 µm for the Visium legacy. Therefore, the 299 × 299 window represents a square tile of 55 µm in dimension for the 10x Visium, and a separate model is trained for this resolution compared to a model trained for a resolution of 100 µm. With this setup, a well-trained model can be applied to histological images (non-ST data) to predict gene expression at the spot level.

First, we focused on breast cancer datasets, including an in-house dataset and multiple publicly available breast cancer datasets that differ in resolution (33 samples from the legacy ST platform and nine samples from the 10x Visium platform) and sample preservation types, with two fresh frozen and one Formalin-Fixed Paraffin-Embedded (FFPE) sample among the nine Visium samples. To benchmark STimage, we compared it with three existing methods: STnet^9^, Hist2Gene^10^, and Hist2ST^11^, using the 33 samples from the legacy ST platform^9^. Model training, evaluation, and testing were conducted using a leave-one-out cross-validation (LOOCV) strategy to benchmark the prediction accuracy of the regression model against the existing models.

To test the generalisability of the model to datasets generated from different laboratories and independent cohorts, we used the nine-sample Visium dataset, including six Visium samples for HER2+ breast cancer^14^ and three from the 10x Genomics public dataset. The dataset was divided into 70% training data and 30% validation data. The test data included one FFPE sample from 10x Genomics and one fresh frozen sample from the HER2+ dataset^14^. For independent validation, we randomly selected three whole-slide histological images from the TCGA HER2+ cohort. In this way, the trained models from the STimage software on the Visium dataset were applied to assess their applicability to non-spatial data.

We also produced subcellular-resolution data using Xenium for five samples, and we demonstrated the unprecedented capability of using this data to validate gene expression predictions as an out-of-distribution (OOD) test. We then further extended the model to predict cell types for hundreds of thousands of individual cells. In all model training scenarios, the data were split at the sample level (WSI), not at the tile level, to avoid overfitting. We also performed OOD evaluations in several different scenarios. The regression model trained on the Visium breast cancer dataset was used to make predictions on H&E images from the legacy ST HER2+ dataset^9^, to evaluate cross-platform prediction, and on H&E images from in-house CODEX spatial proteomics data of skin cancer, to compare the predicted gene expression with the protein modality.

Beyond testing for breast cancer data, we also assessed STimage on two other cancer types (skin and kidney cancer datasets) and a non-cancer liver immune disease dataset (primary sclerosing cholangitis). The in-house skin cancer dataset includes 13 melanoma patient samples, processed using the Visium FFPE protocol. The kidney cancer dataset comprises six Visium samples^15^, and the liver dataset includes four Visium samples^16^. LOOCV was used to evaluate STimage model performance for each dataset separately.

### 2.2 Preprocessing of input image

Technical variation between H&E images can be caused by various factors, such as different staining procedures, imaging hardware and settings, or operators. Such technical artefacts can affect model training and testing, negatively impacting generalisability. In STimage, we perform stain normalisation for each image, such that the mean R, G, and B channel intensities of the normalised images are similar to those of the template images, while preserving the original colour distribution patterns. STimage uses StainTool V2.1.3 to perform Vahadane^17^ normalisation as the default option. We show that this processing step improves model performance.

In addition, the nature of tissue sectioning will inevitably result in some tiles containing low tissue coverage. These tiles should be removed. STimage uses OpenCV2 for tissue masking and removes tiles with tissue coverage lower than 70% (default).

### 2.3 STimage regression model

We utilised both sequencing data and H&E image pixel intensity data to train STimage regression models for predicting spot-level gene expression based solely on imaging data.

We adopted the NB-distributed likelihood **NB r**, **p** for the STimage regression model, where 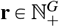denotes the “number of successes”, **p ∈ (**0, 1)*^G^* the “probability of each trial’s success”, and *G* is the number of (assumed) independent genes. The parameters of the NB distribution, **r** and **p**, are estimated from the spatial spot image **s** using an image feature extractor (ResNet50 by default), *f _θ_*, with learnable pre-trained parameters θ from the ImageNet dataset. The functions *h*_ω_ and *h ^η^* represent parameter spaces containing the weights and biases of the neural network layers that connect the latent space 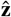 to the two output neurons **r** and **p**.

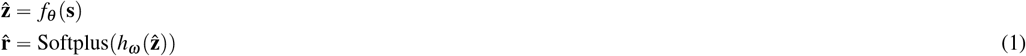

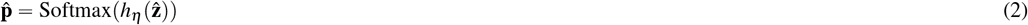

The original STimage regression entails the ResNet50 CNN *f_θ_* mapping a H&E spot tile, **s**, to a latent representation 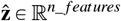, which is then fed into separate output layers (1) and (2), estimating 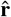 and 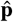, respectively. We denote estimates for the *g*-th gene’s likelihood parameters by 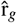 and 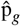. Therefore, the STimage regression model defined *g*-th (random) gene expression *X_g_*| **s**, as a function of H&E spot **s**, by

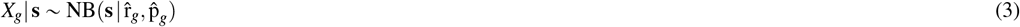

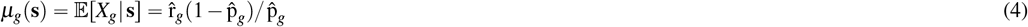

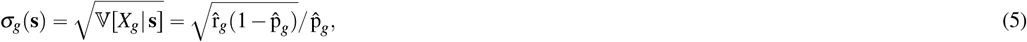

where *µ_g_* **s** in (4) is the mean prediction for the *g*-th gene’s expression with expectation operator 𝔼 [.], and γ*_g_* **s** in (5) is the standard deviation function with variance operator 𝕍 [.].

Overall, the ResNet50 CNN layers are optimized to take H&E images as input to estimate each gene expression’s predictive distribution with NB’s parameters *r_g_* and *p_g_* (for all *g ∈*{ 1*, …, G*}, Eq. (3)). The ResNet50 base, *f_θ_*, is used to perform convolution operations and spatial filtering to extract a three-dimensional feature map (i.e., tensor) from input image tiles. The pretrained *f_θ_* of the ResNet50 includes the GlobalAveragePooling reduction operation^18^ to convert the three-dimensional tensor to the feature vector **z**, whereby each **z** corresponds to a single tile/spot within a H&E image.

The fully-connected output layers *h _ω_* and *h _η_* were fine-tuned by maximising the NB’s negative log-likelihood,

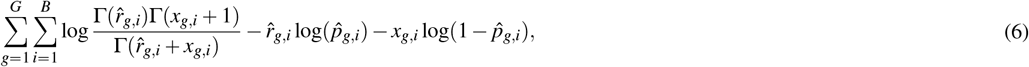

with batch size *B*, Gamma function Г (·), and *x_g,i_* representing the observed RNA count of the *g*-th gene (i.e., gene expression).

To assess the model’s predictive capability, we first trained our model on a panel of 14 marker genes, which included three immune marker genes (CD74, CD63, CD81) and eleven cancer-associated genes (COX6C, TP53, PABPC1, GNAS, B2M, SPARC, HSP90AB1, TFF3, ATP1A1, FASN, VEGFA). Two 10x fresh frozen breast cancer samples were used (replicate 1 for training and replicate 2 for testing) to evaluate the model’s performance in biomarker prediction. We then extended the model to nine breast cancer datasets, initially using the top 1,000 most variable genes and subsequently applying it to a comprehensive set of the most relevant functional genes (1,522 genes). The top 100 genes were selected as the most predictable based on their Pearson’s correlation coefficient (PCC). During training, the model also learned the relationships between these genes.

We further improved STimage by integrating the latest foundation models for image feature extraction. The H&E image accompanying the ST data is preprocessed in the same way as described earlier. Instead of using ResNet50 as the backbone of STimage, we utilise a foundation model to convert H&E tile images into feature embeddings. These embeddings are then connected to STimage’s neural network layers to estimate gene expression distributions. To demonstrate the improvement, we incorporated Virchow2, one of the most advanced foundation models, which has been trained on 3.2 million whole-slide H&E images and contains 632 million parameters. Due to its extensive training dataset and large-scale parameterisation, Virchow2 serves as a more suitable image feature extractor compared to traditional models such as ResNet50. Further, we created an ensemble model named **FSTimage_ens**, where we integrated multiple foundation models into the STimage base model. Each foundation STimage model contains one foundation model as the upstream image extractor and an STimage negative binomial module. The foundation models used are: *GigaPath, H-Optimus, Phikon, cTranspath, CONCH, UNI*, and *Virchow2*. We then aggregated by averaging the results from these models to achieve a final FSTimage ensemble (**FSTimage_ens**) prediction. We found that *STimage, Virchow2_STimage*, and **FSTimage_ens** performed better than any other models that we compared to, including *THItoGene* and *IGI_DL*.

### 2.4 Uncertainty assessment

STimage estimates a NB distribution to model each gene’s (random) expression *X_g_* ∈ {0, 1, · · ·} conditional on the H&E spot **s**, i.e., 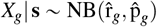. A single STimage model produces the distribution 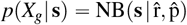. An ensemble of Stimage models produces a distribution *p* (*X_g_* **s**, Θ). Therefore, following the law of total variance, we can obtain two terms accounting for uncertainty:

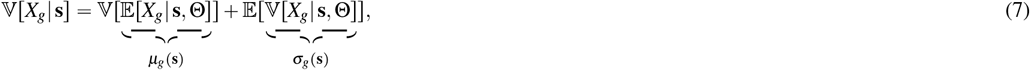

where the variance of the mean estimates, 𝕍 [*µ_g_* (**s)**], defines epistemic uncertainty, and the average of variances, 𝔼 [α*_g_* (**s)**], defines aleatoric uncertainty. Aleatoric uncertainty reflects the noise inherent in the data. Epistemic uncertainty reflects the model’s limitations (e.g. lack of information due to small sample sizes or suboptimal model parameters) and can be reduced with more data and/or improved models.

Applying the formula (7), we quantify total uncertainty for the prediction of each gene in each spot. Briefly, we trained 30 STimage models for approximately 1,522 functional genes separately, using the same training dataset, each with a unique random seed. Corresponding values of *X_g_* were predicted for individual spots **s** using each trained model with model parameters Θ. Based on the 30 individual STimage models, we computed five ensemble models. Each STimage ensemble model was derived by averaging NB estimates *µ _g_*(**s)** from five randomly pooled STimage single models. For each of the single models and ensemble models, PCC was then calculated as the prediction accuracy metric.

### 2.5 Model interpretation of STimage

STimage implements model explainability in order to highlight important segments/features (nuclei) from the image that contribute to the high expression of a gene. We used a local model-agnostic approach (LIME)^19^, in which different segments of the images are repeatedly perturbed to measure how the predictions made by the trained model change, in order to score each individual segment.

Morphological diversity among cells in the tumour is an important hallmark that provides clues for histopathological assessment. In order to obtain a score for each nucleus, we used Cellpose-3 segmentation^20^ to segment the nuclei from H&E images. We performed an image transformation, converting RGB images to Haematoxylin-Eosin-DAB (HED) format before applying the Cellpose segmentation. Inputs required by LIME are a trained regressor/classifier, an image tile, a gene name, and a segmentation function. Using LIME, we identified nuclei segments corresponding to high or low feature importance in the regression model. Given below are the formulas used to compute the LIME values.

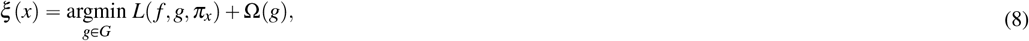

Where

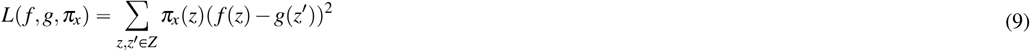

LIME finds a simple model *g* that can approximate our complex neural network model *f*. The total loss in LIME is given in Eq. (8). The term *L*(*f, g, π_x_*) in (8) is a weighted sum of squared errors (SSE), where the weights are given by π & *z*, depending on the proximity of the local neighbourhood data point *z* to *x*. This ensures that points close to *x* are given greater weight.

Eq. (9) creates a local approximation of our trained model in the neighbourhood of a given input image *x* using a simple, interpretable regression model *g*, where *G* is a set of sparse linear regression models and *π_x_* denotes the kernel function that defines the local neighbourhood around data point *x*. Here, *f* (*z*) is the prediction made by the neural network for features *z*, and *g* is a simple model that approximates the behaviour of *f* in the locality of *x*, as defined by π*_x_*. The vector *z*^’^ is an explainable subset of *z*. The second term in (8) is a regularisation term applied to *g*, which assigns zero weight to non-important input features.

#### STimage Web Application

To provide an interactive, visual interpretability option, we have developed the STimage Web App, where users can select image tiles and genes of interest to visualise the nuclei important for the prediction of gene expression. The Web App allows users to either use a pre-trained model or upload their own trained model for interpretability visualisation. We provide trained STimage regression models for breast cancer, kidney cancer, liver disease, and skin cancer. The Web App can be accessed at: https://gml-stimage-web-app.streamlit.app/.

### 2.6 STimage classification model

We reasoned that categorical prediction of gene expression—mapping whether genes are present or absent, or highly or lowly expressed in specific tissue regions—could already have significant clinical potential.

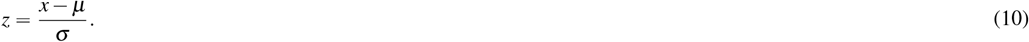

We transformed the raw gene expression values (continuous) to *z*-scores using (10), and categorised them as “Low” for *z*-scores below zero and “High” for *z*-scores above zero. For the classification model, we used a pre-trained ResNet50 (optionally with a fine-tuning layer) as a feature extractor for all tiles, and then applied a regularised logistic regression classifier, optimising the log-loss (binary cross-entropy) function.

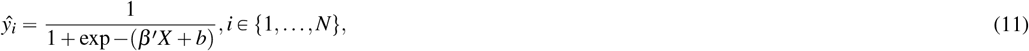

We used the “elastic net”^21^ to balance model performance and complexity by applying both L1 and L2 penalties. In Eq. (11), the coefficient set *β*=(*β*_1_, …, *β_n_*) and ***X =(****x*_1_*, …, x_n_)* denote the latent space features (*N* 2048 features).

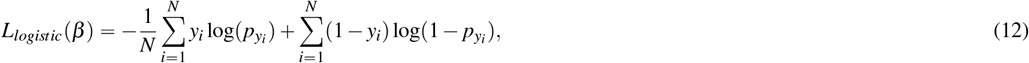

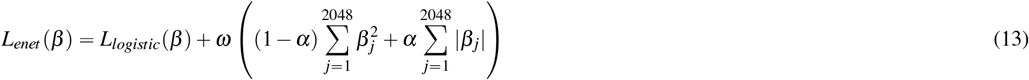

The elastic loss *L_enet_* in Eq. (13) is the sum of the binary cross-entropy loss (log loss) from Eq. (12) and the regularisation terms.

Here, *α* in Eq. (12) is a mixing parameter defining the relative contribution of RIDGE and LASSO to the penalty term. For α=1, the ElasticNet function reduces to LASSO regularisation (~ logistic loss + sum of absolute values of weights), while for α*=* 0, the function reduces to RIDGE regularisation (~ logistic loss + sum of squares of weights). ω is the regularisation parameter, and we set the regularisation strength to ω= 0.1 (a smaller value implies weaker regularisation). The solver “SAGA”^22^ was used for optimisation, as it is faster for large datasets.

The classification model was trained on seven breast cancer samples for the top 100 highly predictable breast cancer genes (from the STimage regression model) and tested on 10x FFPE and fresh frozen HER2+ (1160920F) samples.

### 2.7 Benchmarking and performance assessment

We applied a LOOCV strategy on 33 samples (from a total of 36, with 3 excluded due to low quality) from the HER2+ ST dataset to evaluate STimage regression model performance and benchmark it against existing methods such as STnet^9^, HisToGene^10^, and Hist2ST^11^. Briefly, all models were evaluated using the same data-splitting LOOCV strategy, where models were trained on 32 samples, and the spatial gene expression was predicted on the held-out sample.

For each gene, Pearson’s correlation coefficient (PCC) was calculated between the predicted and experimentally measured spatial gene expression values. PCC was used as the primary performance metric to assess prediction accuracy. We also computed the autocorrelation Moran’s I index as an additional metric—a weighted correlation measure that incorporates the local spatial context.

To assess generalisation, we evaluated model performance on the top 1,000 highly variable genes (HVGs). After removing genes expressed in fewer than 10% of total spots, 785 HVGs were retained for further analysis. Each gene received a score from applying the model to a patient, resulting in 785 scores per patient for comparison across models.

To ensure fair comparisons, both the HisToGene^10^ and Hist2ST^11^ models were run using the authors’ recommended hyperparameters. However, STnet failed to execute due to technical issues in the original source code, so we re-implemented the STnet model according to the original paper^9^. Once all predictions from all models were collected, we obtained 103,620 values (33 models × 4 methods × 785 genes) for comparative analysis.

We also benchmarked STimage against the other methods using an out-of-distribution (OOD) test dataset. In this scenario, all models were trained on two fresh frozen breast cancer tissue samples from 10x Genomics and tested on an FFPE tissue sample, which we considered as an OOD sample due to the substantial differences in data generation protocols between these two sample types—namely, poly-A capture for fresh frozen versus probe-capture for FFPE protocols. Using the same gene filtering approach as described above, we obtained 982 of the top 1,000 HVGs in both the training and prediction datasets.

Hist2ST and HisToGene include a vision transformer (ViT) component as part of their neural networks, which requires loading all spot tile images and expression values from a single tissue into the model in one batch. Consequently, running these models requires greater computational resources (particularly memory) when applied to Visium data, which contain approximately 10 times more data than legacy ST datasets. Training of Hist2ST and HisToGene failed on a computing cluster equipped with an NVIDIA Tesla V100 SXM2 32 GB GPU, necessitating a workaround: we split the Visium tissue into multiple smaller subsets using a sliding window approach to reduce memory load.

Each subset contained a square tissue region of 25 × 25 spots with no overlap. Each subset was treated as an individual dataset, which reduced the number of spots loaded per batch while preserving the spatial neighbourhood structure. All subsets from the two fresh frozen samples were used to train Hist2ST and HisToGene models using their default hyperparameters. The trained models were then applied to predict gene expression in the FFPE dataset. Model performance was assessed using both Pearson’s correlation coefficient (PCC) and Moran’s I index.

### 2.8 A new geometric metric to better assess model predictions of spatial patterns

The commonly used metric in gene expression prediction tasks, PCC, is known to be influenced by outlier noise and is unable to capture spatial patterns. To address this, we developed a new metric for evaluating model performance in terms of how well predictions match ground truth spatially. For a given gene, we standardise the predicted and ground truth values to *z*-scores and then categorise them as binary values, based on whether the values are greater or less than zero. These binary values are then organised into a spatial matrix, creating a binary image where gene expression is plotted according to spot coordinates. To minimise noise and retain reliable prediction patterns, we apply Gaussian filtering with a sigma value of 1 to both rows (spatial *X*) and columns (spatial *Y*). After obtaining the refined binarised prediction and ground truth images, we compute the Intersection over Union (IoU) scores to assess the overlap between the spatial patterns of the ground truth and the prediction. This score effectively reflects the accuracy of predicting spatial expression patterns.

### 2.9 Cell type classification model for Xenium single-cell ST data

We modified the Hover-Net model^23^, a CNN-based instance classification and segmentation model, to enable cell segmentation and cell type prediction for Xenium spatial transcriptomics data at single-cell resolution. First, we generated cell type labels by annotating the cells in the Xenium data with individual cell types using Seurat reference-based integration with the human breast cancer atlas from^14^. We merged similar cell type labels into four categories: immune cells, stromal cells, cancer epithelial cells, and endothelial cells. The Xenium labels were then mapped to the H&E image by applying Scale-Invariant Feature Transform (SIFT) registration^24^ between the Xenium DAPI channel and the H&E image. This enabled us to obtain, for each tissue sample, segmentation labels with unique cell types for nearly every cell in the H&E images (i.e., 1,294,600 cells).

To generate the training and test data, the H&E images were divided into 256 × 256-pixel tiles, and tiles containing fewer than two nuclei were filtered out. The model was trained using four of the five breast cancer tissue samples and evaluated on one held-out test sample.We also calculated the mean IoU score for each LOOCV test sample. We compared our model with the recent CellViT model ^25^, a vision transformer-based model that was trained to predict single-cell segmentation and cell type classification.

### 2.10 Validation using non-spatial data and assessing predictive values for survival and drug responses

To perform external validation, assess robustness, and demonstrate the clinical applicability of STimage, we applied the model to the TCGA dataset. For this analysis, H&E images were downloaded from TCIA^26^, and bulk gene expression and clinical data for corresponding patients were downloaded from https://www.cancer.gov/ccg/research/genome-sequencing/tcga. We obtained 3,066 images from TCIA. A total of 1,532 images without matched bulk and clinical data were excluded from the analysis, and the remaining 1,534 images were processed for quality control using HistoQC. 285 images failed HistoQC input step as they were unreadable. For the remaining 1,249 images, we filtered out low quality images using the following parameters: blurry_removed_percent ⩾ 0.9, small_tissue_removed_num_regions ⩾ 97.5^th^ percentile, and pixels_to_use <5^th^ percentile. These thresholds were chosen based on the empirical distribution of quality parameters, removing extreme values. Using these filters, 215 images were excluded. Only images that passed all the filtering criteria were then used for further analysis. We finally used 1,034 breast cancer images from 670 patients and predicted the gene expression of 1,522 predictable genes. Predicted gene expression for each image was averaged across all spots to compare with the ground truth TCGA bulk gene expression. We measured the performance of STimage by calculating Pearson Correlation (PCC) between the average predicted expression across all the spots with the true bulk gene expression for the matched samples. We also computed the performance of STimage for predicting top-300 predictable genes for three subtypes (HER2+, Lumnial and TNBC). Patient subtypes were defined according to the criteria described in^27^.

STimage predictions were further used to perform survival analysis and stratify patients into low- or high-risk groups. We first built univariate Cox regression models for the top 300 predictable genes using the function ‘coxph’ from the R package `survival`^28^. Cox regression models were built separately for true and predicted gene expression. Days to death and days to last follow-up (for patients still alive) were used as survival times, and survival status (alive/dead) was used as the event indicator. The univariate models were evaluated using the concordance index (c-index) for both predicted and true survival outcomes.

The top five genes with the highest c-index were used to build multivariate Cox regression models. We used a 3-fold cross validation strategy with 100 repetitions to evaluate the multivariate models.The median survival risk score from the multivariate model was used to stratify patients into low- or high-risk groups. Univariate and multivariate models were constructed separately for each subtype. For both bulk and predicted data, the average c-index for the multivariate models was found to be over 0.75 across all subtypes. Model performance was assessed using the ‘concordance’ function from the `survival`package. Univariate Cox regression models and Kaplan–Meier survival curves were computed using the `survival`package. Multivariate models were constructed using the R package `ClassifyR`^29^.

To further assess the clinical utilities, we added to our previous survival analysis an independent assessment of the ability to predict responses to drugs. Here we analysed a cohort of 168 breast cancer patients treated with neoadjuvant and chemotherapy with (n = 65) or without HER2-targeted therapy^30^. 161 patients were assessed at surgery using the residual cancer burden (RCB) classification. 42 (26%) had a complete response (pCR), 25 (16%) had a good response, 65 (40%) had a moderate response and 29 (18%) had extensive residual disease. We evaluated the performance of classification models using true or predicted gene expression for classifying pCR vs RD using a an RBF-kernel support-vector machine. We used 1139 genes in the functional gene list that were present in the data. The continuous predicted gene expression data was log-transformed, while the raw count gene expression data was CPM-normalised and standardized. We applied a nested 5-fold stratified cross-validation: each outer fold used four folds for training and one for testing, with an inner 5-fold split for model selection. This process ensured every patient was tested once. AUC was calculated for each fold, and the average AUC was reported as the final performance metric.

## 3 Results and Discussion

### 3.1 STimage model - distribution prediction, ensemble, interpretability and uncertainty estimates

First, in the STimage pipeline, we implement an important—yet often underestimated—image preprocessing step (Figure 1a). STimage performs stain normalisation to account for technical variation between image inputs used during model training. To remove uninformative tiles, we calculate tissue coverage and filter out tiles with low coverage. In STimage, image tiles are either used to fine-tune the ResNet50 model (as a minibatch of 64 tiles by default) or passed through pretrained foundation models to be transformed into feature embeddings (Figure 1b). All latent features are then connected to a layer of neurons representing the parameters of the negative binomial (NB) distribution for each gene, which is optimised using a log-likelihood loss function. Alongside the distribution estimates, we also quantify model prediction uncertainty (Figure 1b). This is a critical feature not available in existing tools, yet essential given the high variability expected in cancer datasets. Multiple genes can be trained simultaneously, with each gene corresponding to a pair of NB parameters. By sampling from the NB distribution, STimage can learn to predict the expression of individual genes or gene groups. Importantly, we incorporate model interpretability via LIME (Figure 1b).

**Figure 1.**
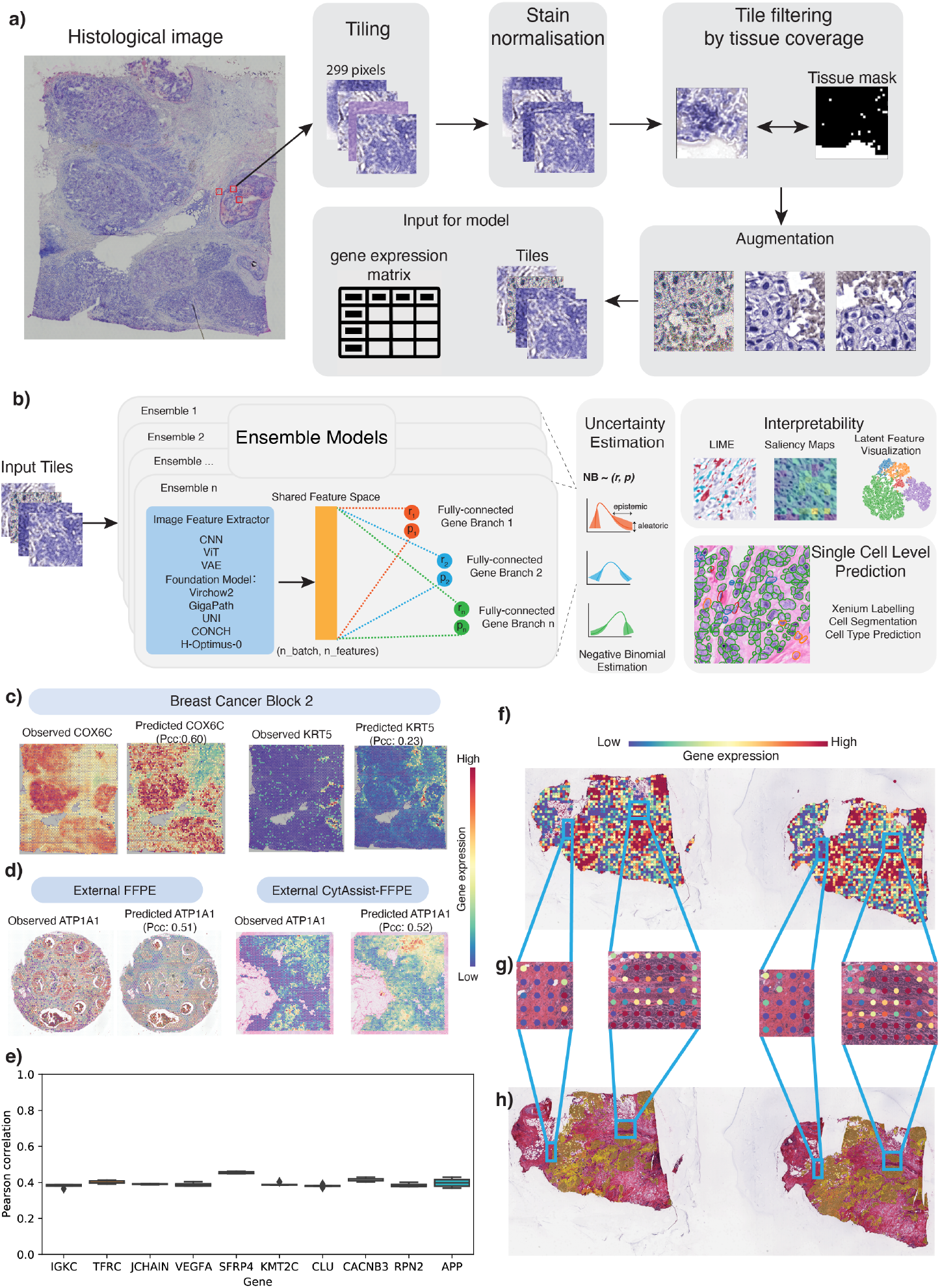
Overview of the robust and interpretable STimage model. **a**, STimage image preprocessing workflow includes image tiling, stain normalisation, quality control (QC) for tiles, and image augmentation. **b**, STimage CNN–NB interpretable regression model. The STimage model consists of an image feature extractor (CNN, ViT, or foundation models) and negative binomial (NB) layers for estimating gene expression distributions. Negative log-likelihood is used as the loss function. **c**, ST-measured (observed) and STimage-predicted gene expression for the highly abundant cancer marker COX6C and the lowly abundant but spatially distinct keratin marker KRT5 on the test dataset. PCC between predicted values and observed values across all spots (measured by spatial transcriptomics) is shown. **d**, Model performance for an out-of-distribution FFPE dataset from 10x Genomics. **e**, PCC values for the 10 most predictable functional genes (i.e., top 10 PCC scores out of 1,522 functional genes), calculated for two external FFPE datasets using an ensemble model. **f, g, h**, STimage regression model applied to non-spatial whole-slide images (WSIs) of a HER2+ patient from the TCGA dataset. Two adjacent tissue sections were used to assess reproducibility. **f**, The WSI was cropped into non-overlapping tiles of size 299 × 299 pixels. A model trained on nine Visium samples was applied to predict expression of the cancer marker gene COX6C. Each tile is treated as a spatial spot; red and blue colours indicate high and low predicted COX6C expression, respectively. **g**, Two zoom-in views of tumour-enriched areas. Predicted cancer marker expression is highly correlated with tumour morphology and consistent across replicates. **h**, Pathologist annotation of tumour regions (indicated in brown) in the two replicate tissue sections.

Several existing programs for gene expression prediction—such as STnet^9^, Hist2ST^11^, Hist2Gene^10^, DeepSpace^12^, and Xfuse^31^—produce fixed-point estimates without any uncertainty quantification. In contrast, STimage introduces an approach to quantify and mitigate uncertainties in gene expression predictions, distinguishing between uncertainty due to model knowledge gaps (epistemic) and inherent variability in the data (aleatoric) (Figure 1b).

### 3.2 New prediction performance metric and STimage accuracy

To assess model performance, we first visually compared the prediction of an abundant cancer marker (COX6C), broadly expressed across the tissue, with a less abundant marker, KRT5, expressed specifically in myoepithelial cells surrounding Ductal Carcinoma In Situ (DCIS) regions. The model trained on nine Visium breast cancer datasets was used to predict expression in a new Visium test dataset, showing highly consistent spatial patterns between predicted and observed values (Figure 1c). Quantitatively, using the spatial autocorrelation metric Moran’s I, we observed a strong positive correlation (i.e., high-high or low-low clustering) for COX6C expression across most spatial spots (Figure S2a). We applied the same model—trained on the nine fresh frozen Visium samples—to two external, previously unseen out-of-distribution FFPE samples: one measured using the version-1 FFPE protocol and the other with the FFPE CytAssist protocol (Figure 1d). Results for one representative cancer-immune marker, ATP1A1, are shown for both FFPE test samples (Figure 1d). Results for all 14 cancer-immune markers, evaluated using PCC and Moran’s I metrics, demonstrate consistently positive scores, although performance varies across samples (Figure S2c,d,e). This spatially-aware quantification offers a distinct advantage over non-spatial models such as

HE2RNA^7^, which do not provide per-tile gene expression predictions.

### 3.3 Prediction performance — more robust across diverse datasets and sample types

To assess the robustness of the approach, we increased heterogeneity in the training and testing data by including samples from different protocols, laboratories, and populations. We trained a model using nine Visium datasets from fresh frozen tissues, with three datasets generated by 10x Genomics and six by Swarbrick’s Laboratory^14^. Using an ensemble approach, the model trained on fresh frozen samples was applied to two external FFPE samples processed using Visium protocols. The top predicted gene, ATP1A1, showed high PCC values (0.51 and 0.52) and consistent gene expression patterns (Figure 1d). Assessing model performance across 1,522 functional genes (trained simultaneously), we found that predictions were positively correlated with measured values for 1,136 of 1,522 genes. The top 10 genes achieved PCC scores around 0.4 (Figure 1e). Notably, model performance was consistently higher when our image normalisation strategies were applied (Figure S3). We also thoroughly assessed the impact of image tile resolution and found that the default 299 × 299 tile size in STimage yielded the highest predictive performance (Figure S4). These findings highlight options for improving robustness.

The trained model incorporated both the public dataset from 10x Genomics and Swarbrick’s dataset, while testing was performed on an independent dataset generated by a European laboratory, consisting of 33 tissue samples. STimage was benchmarked against six other approaches: five software tools—THItoGene, IGI_DL, STnet, His2gene, and Hist2ST—and an alternative model, ViT_NB. STimage consistently outperformed all other methods across the majority of test samples, except for six (out of 33), where its performance was comparable to the top-performing models (Figure 2e). To further enhance performance, we developed an ensemble model, FSTimage, which integrates multiple foundation-STimage models. Each model combines a foundation model as the upstream image extractor with the STimage negative binomial module. The foundation models used include GigaPath^33^, H-Optimus^34^, Virchow2^35^, Phikon^36^, CTransPath^37^, CONCH^38^, UNI^39^, and Virchow^40^. Final FSTimage predictions were obtained by averaging results across these models. Our findings indicate that STimage, Virchow2_STimage, and FSTimage consistently outperformed all other models compared.

**Figure 2.**
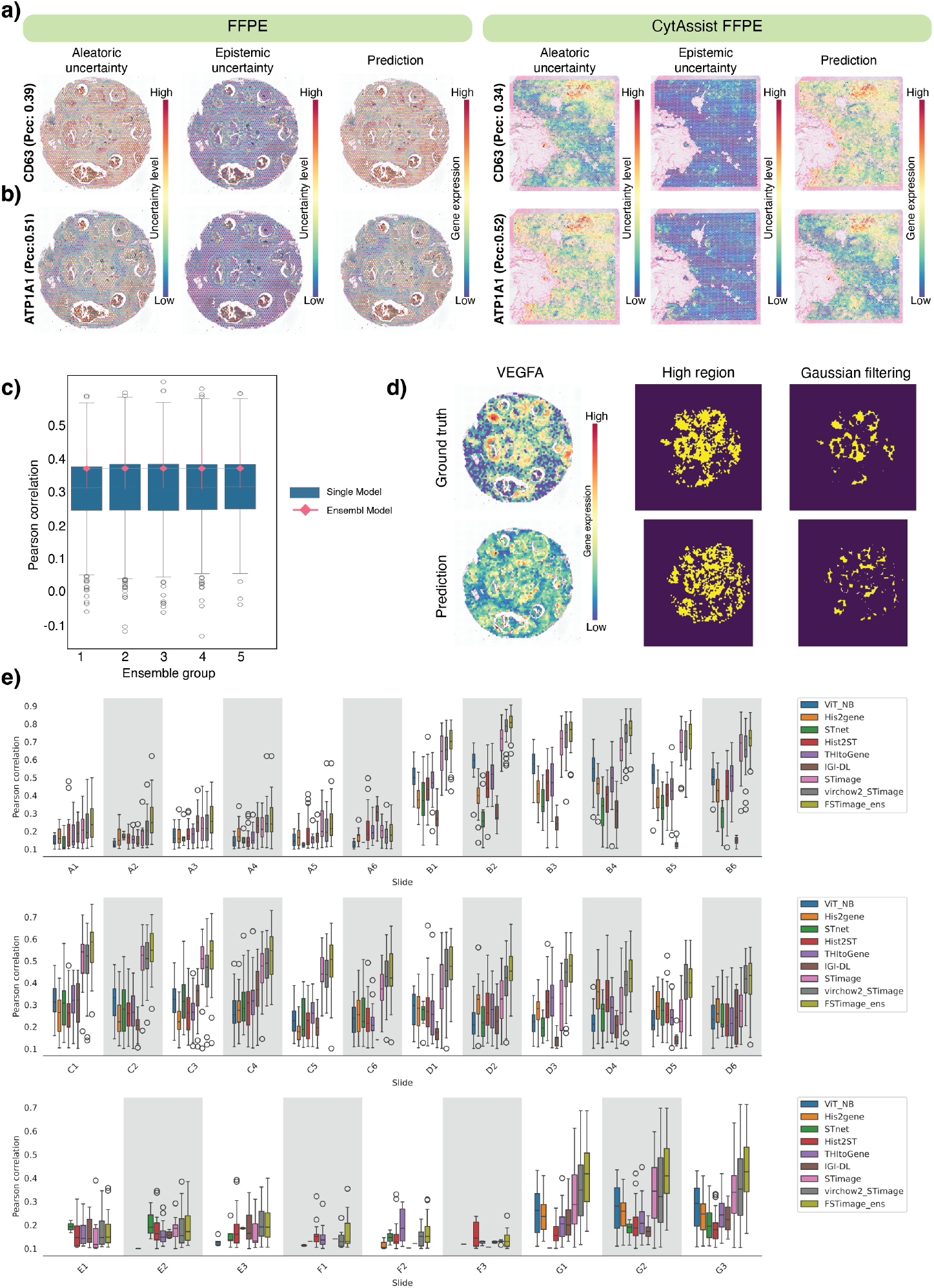
Distribution-based estimation and benchmarking of the STimage CNN–NB model. **a**, Uncertainty estimation and predicted gene expression for CD63 in two external FFPE datasets. The red–blue colour gradient indicates high to low uncertainty levels. **b**, Uncertainty estimation and predicted gene expression for the cancer gene ATP1A1 (the top predicted gene with high PCC). **c**, Performance of five ensemble models compared to 10 single models within each ensemble group. The x-axis shows five ensemble groups, each containing 10 runs. The y-axis shows the PCC between predicted and ground truth gene expression. Boxplots show the distribution of prediction performance across 10 single models (with different random seeds) for two tissue slides per group. The connected lines represent ensemble models (aggregating all single models per group), with error bars indicating standard deviation across genes. **d**, Customised IoU score developed to better evaluate spatial model performance. **e**, Comprehensive benchmarking against STnet, HisToGene, THItoGene, IGI-DL, and Hist2ST using 33 tissue slides from the HER2ST dataset^32^. The plot shows PCC values between predicted gene expression of HVGs and ground truth measurements. Each boxplot represents one test sample in the Leave-One-Out Cross-Validation (LOOCV) experiment per method. ‘virchow2_STimage’ refers to the STimage model using foundation model embeddings from Virchow2. ‘FSTimage_ens’ denotes the ensemble model using five foundation model weights from HEST.

To further evaluate robustness and increase sample diversity, we incorporated additional datasets, including kidney cancer (fresh frozen, *n* = 6, 15,632 spots), non-cancer liver disease (FFPE, *n* = 4, 19,967 spots), and skin cancer (FFPE, *n* = 10, 14,405 spots) (Figure S7, S6, S5). We used a leave-one-out model training and testing strategy to evaluate model performance on individual patient samples. Among the top 100 most predictable genes, GO enrichment analysis revealed terms corresponding to relevant tissue biology across the three GO databases (Figure S7a–c, S6a–c, S5a–c). The PCC between predicted and ground truth gene expression values among the top 100 genes showed a positive correlation, ranging from 0.2 to 0.8 (Figure S7d, S6d, S5d). Performance also varied across patients, suggesting tissue heterogeneity. Representative spatial expression plots are shown in Figure S7e–g, S6e–g, S5e–g.

### 3.4 Prediction uncertainty can be quantified for each gene

In addition to improving robustness, ensemble models allow us to quantify uncertainty arising from both the data (i.e., aleatoric uncertainty) and the model itself (i.e., epistemic uncertainty). Existing methods such as STnet^9^ and HisToGene^10^ use a CNN or ViT as an image feature extractor. Both models rely on multi-layer neural networks to predict fixed gene expression values, without estimating variability in their predictions. Hist2ST^11^ employs a zero-inflated negative binomial distribution to predict gene expression distributions, but it does not account for epistemic uncertainty. To evaluate these new capabilities in STimage, we used the model trained on the nine Visium fresh frozen breast cancer datasets to test its performance on two independent, unseen FFPE tissues from separate cohorts (Figure 2a–d and S1). Two representative genes were selected for comparison: ATP1A1, a breast cancer marker, and CD63, an immune marker. For each gene, aleatoric uncertainty, epistemic uncertainty, and predicted gene expression were visualised in spatial tissue plots (Figure 2a,b). Aleatoric uncertainty (i.e., the average of predicted NB variance values across single models) was found to be higher than epistemic uncertainty (i.e., the variance of NB mean estimates) in both FFPE samples. As expected, both uncertainty types showed a positive correlation with gene expression levels. Interestingly, lower epistemic uncertainty often coincided with higher aleatoric uncertainty, as observed for ATP1A1 (Figure 2b), thereby increasing confidence in the predicted NB parameters (i.e., mean and variance).

In addition to lower epistemic uncertainty, ATP1A1 also showed higher predictive performance (PCC: 0.52; Figure 2b) compared to CD63 (PCC: 0.34; Figure 2a). This finding aligns with interpretations of epistemic uncertainty, which reflects the confidence in model predictions^41^. In this case, the higher epistemic uncertainty observed for CD63 may be partially attributed to the fact that the test samples were generated using different technological platforms compared to those in the training data. As a result, the input data were considered out-of-distribution.

### 3.5 An ensemble approach improves uncertainty performance

To increase model robustness across a broad range of diverse, unseen datasets, we trained an ensemble of STimage regression models. Ensemble models have been shown to improve generalisation in the transcriptomics context^42^. We observed variation in model predictions, even when using the same training dataset and model architecture but with different random seeds. To assess and mitigate this variability, we analysed the distribution of performance across individual models within 12 different groups and for two distinct tissue slides (a total of 30 models). The results are shown as box plots in Figure 2c. Each box represents the average PCC from independent predictions made by 10 single models for each of 12 genes (out of a total of 14 trained genes; 2 were not presented in the additional CytAssist FFPE dataset). The lines represent ensemble models obtained by averaging predictions from all 10 single models within each group, with error bars indicating the standard deviation across different genes (12 genes). Although there is considerable variation among individual models, their performance remains consistent across ensemble groups. Furthermore, the ensemble models outperform the single models and exhibit reduced variability across all groups (Figure 2c). We further demonstrate that this ensemble strategy also applies to the FSTimage_ens model, where embeddings from different foundation models are combined to further enhance performance (Figure 2e).

### 3.6 Out-of-distribution performance for models trained on a small dataset

To assess model performance on out-of-distribution (OOD) data, we designed an experiment where the STimage model and three other models—STnet, HisToGene, and Hist2ST—were trained on fresh frozen samples (Visium poly-A captured data) and tested on FFPE data (Visium probe-hybridisation data). The probe-hybridisation data detected more genes and provided more information per gene. As the two datasets were derived from different tissue types and technologies, they were considered OOD (Figure S1).

Previous studies, such as Hist2Gene^10^ and Hist2ST^11^, showed limited performance when trained on small datasets. In contrast, we found that the STimage model with a fine-tuning option improved performance on small training sets compared with models using ViT for feature extraction. For a robust comparison, we predicted expression levels for the top 1,000 highly variable genes (HVGs) and computed both PCC and Moran’s I scores for each gene. Across all 1,000 genes, STimage consistently outperformed the other three models.

We further evaluated robustness by testing the prediction performance for each of the 1,000 HVGs, comparing STimage to the three other methods (Figure SS1). Notably, methods based on ViT (e.g., HisToGene and Hist2ST), which generate attention maps across tissue sections, require substantially more memory. These models become challenging to run on datasets with more spatial measurements—a scenario increasingly common with recent advances in spatial transcriptomics. STimage uses a CNN-based image extractor that treats each spatial measurement as an independent data point, enabling scalability to datasets with high spatial resolution. Additionally, STimage leverages pre-trained models for feature extraction, an approach not used by ViT-based methods. We further evaluated robustness and increased sample diversity by incorporating additional datasets, including kidney cancer (fresh frozen, *n=* 6, 15,632 spots), non-cancer liver disease (FFPE, *n=* 4, 19,967 spots), and skin cancer (FFPE, *n=* 10, 14,405 spots) (Figure S7, S6, S5). We used a leave-one-out training and testing strategy on tissue samples to evaluate model performance for each patient. Among the top 100 most predictable genes, GO enrichment analysis revealed terms consistent with the underlying tissue biology across the three GO databases (Figure S7a–c, S6a–c, S5a–c).

The PCC between predicted and ground truth gene expression values among the top 100 genes ranged from 0.2 to 0.8 (Figure S7d, S6d, S5d), demonstrating a positive correlation. Performance also varied among patients, highlighting tissue heterogeneity. Example spatial gene expression plots are shown in Figure S7e–g, S6e–g, S5e–g.

### 3.7 Out-of-distribution performance on independent spatial transcriptomics datasets

In the following sections, we further evaluated the robustness of our model by testing it under more challenging scenarios, involving independent datasets from different cancer types (skin and breast cancer), diverse data types, and various experimental protocols, including low-resolution legacy ST, Visium, single-cell resolution Xenium, non-spatial TCGA images, and CODEX data.

The model, initially trained on nine Visium breast cancer datasets using a functional gene list of 1,522 genes, was applied to the HER2ST^32^ legacy ST dataset. In this experiment, H&E images from the HER2ST dataset were tiled at the spatial location of each spot, using the Visium spot size of 55 µm, which is smaller than the original legacy spot size of 100 µm. Gene expression predictions were generated for these smaller tiles, visualised, and evaluated against the ground truth measurements. Figure 3a displays one of the top predictable genes, ESR1, while additional examples—ERBB2 and B2M—are presented in Figure S8. Figure S8a and b show samples with low and high prediction performance, respectively, while Figure S8c provides pathological annotations for these samples. Even in low-performance predictions, we observed consistent spatial expression patterns when compared with both ground truth and pathological annotation. A quantitative assessment of a large number of genes in the HER2ST dataset is presented in Figure S8d. This includes PCC values for 1,146 predicted genes, derived by intersecting the 1,522 predictable genes with the gene list available in the legacy ST data. We found that a model trained on the high-resolution Visium dataset was able to effectively predict data from the legacy Visium protocol. Prediction performance was highest for samples B1–B6 (median PCC values ranging from 0.4 to 0.6), moderate for samples C1–D6, G1–G3, and H1–H3 (median PCC values from 0.2 to 0.4), and lowest for samples A1–A6 and E1–F3. Notably, samples A1–A6 and E1–F3 consistently showed low prediction performance, even in in-distribution datasets, as shown in Figure 2e.

**Figure 3.**
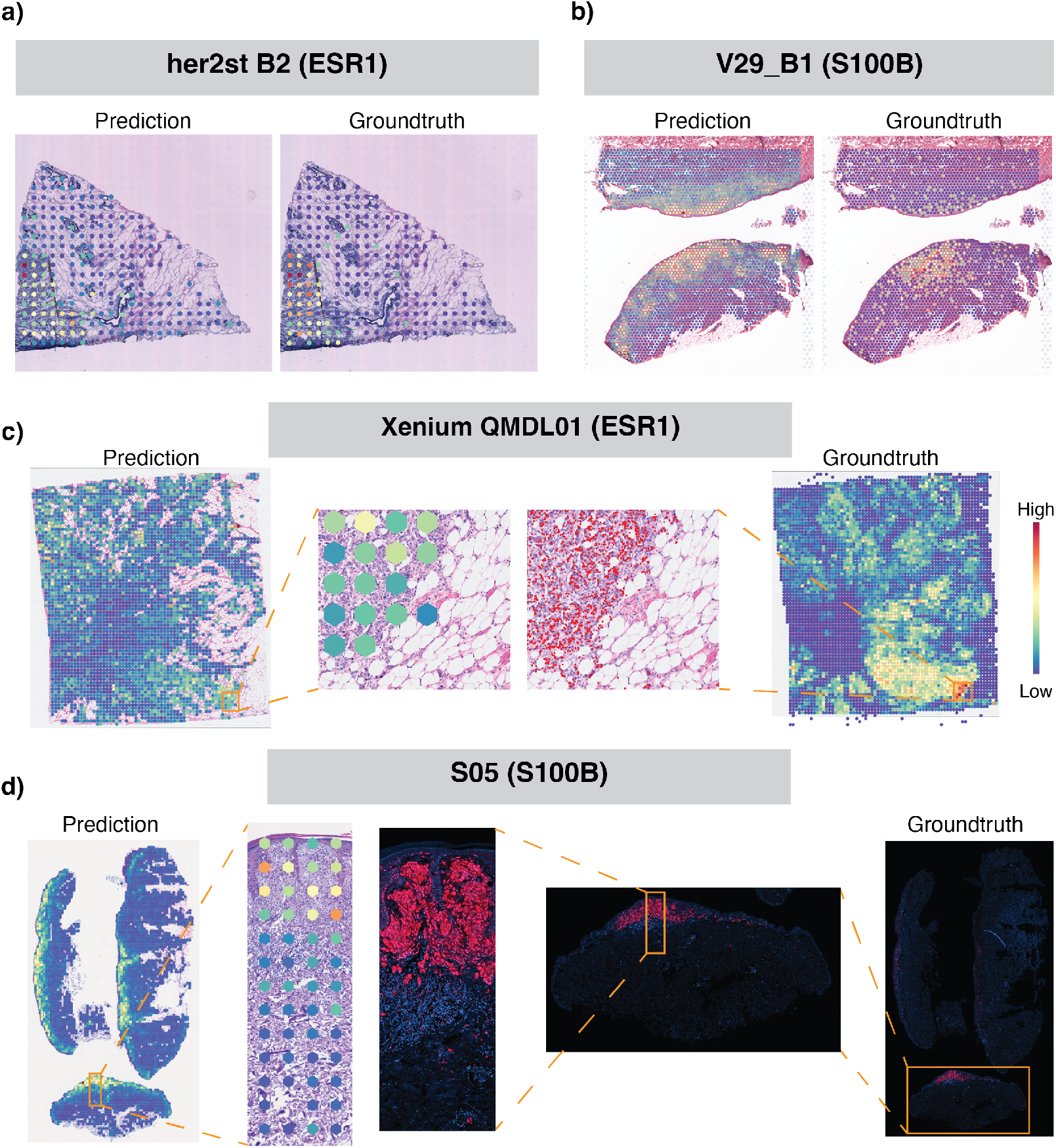
Out-of-distribution assessment of model training on Visium breast cancer data and a functional gene list of 1522 genes, with predictions across various datasets and cancer types using different spatial technologies. **a**, Spatial plot of predicted and ground truth gene measurements of the breast cancer marker gene ESR1 from legacy ST of breast cancer. **b**, Spatial plot comparing predicted and ground truth gene measurements of the skin cancer marker gene S100B from Visium skin cancer data. **c**, Spatial plot of predicted measurements of the breast cancer marker gene ESR1 from the Xenium dataset; the middle section shows a zoomed-in comparison of spot-level predictions and single-cell resolution gene measurements. **d**, Spatial plot of predicted measurements and ground truth protein measurements of the skin cancer marker gene S100B from CODEX skin cancer data; the middle section shows a zoomed-in region showing predicted gene expression at spot level alongside a staining image of S100B protein expression.

In contrast to the previous OOD analysis using a reduced-resolution test dataset (Visium legacy), here we applied the model to higher-resolution data from Xenium. The same model, trained on Visium H&E images, was applied to images from the single-cell resolution Xenium dataset (Figure 3c and S9). Subcellular gene expression measured in the Xenium breast cancer dataset was grouped to match the Visium spot size, serving as ground truth for comparison with model predictions. We again observed highly consistent spatial expression patterns for gene markers such as ESR1 (Figure 3c), although the quantitative correlation was lower than for datasets of the same resolution (Figure S9).

### 3.8 Out-of-distribution performance on another cancer type assessed by spatial proteomics datasets

The same model, trained on nine Visium breast cancer datasets and a functional gene list of 1,522 genes, was applied to H&E images from Visium skin cancer data (Figure 3b and S10), i.e., for a different cancer type. For the first time, we demonstrate that a model trained on breast cancer can predict gene marker expression in skin cancer. Predictions for key melanoma markers such as SOX10 and CD34 were highly consistent with protein data measured by CODEX (Figure S10). The subcellular protein staining signal from the CODEX skin cancer dataset was grouped to match Visium spot size, serving as ground truth for comparison with predictions from H&E images on the same slide (Figure 3d, S10). The ability to predict expression patterns that match protein markers highlights the model’s promise for diagnostic applications.

### 3.9 STimage performs well when trained and tested on other cancer types

Using the same model architecture, we trained the STimage regression model on a skin cancer dataset (Figure S5), a kidney cancer dataset (Figure S7), and a non-cancer liver immune disease dataset (Figure S6). A Leave-One-Out Cross Validation (LOOCV) strategy was used to assess model performance. To further evaluate the biological applicability of the gene expression prediction model on HE images, we performed clustering on the predicted gene expression data to analyse tissue heterogeneity. Overall, the clustering results identify the structural organisation of skin layers (Figure S17a) with expected gene markers. Notably, Cluster 1 corresponds to the melanoma region. We further utilised a set of marker genes associated with epidermal genes, dermal fibroblast genes, and sweat gland genes to characterise clusters. We found that epidermal gene markers were enriched in Cluster 0, while sweat gland genes were enriched in Cluster 5 (Figure S17b). The spatial patterns of the clusters generally align with the HE morphology. Overall, STimage performed consistently well, and the top predictable genes were highly relevant to cancer biology and the tissue of origin.

### 3.10 Classification model provides complementary information on tissue regions with high vs low gene expression

In addition to predicting continuous gene expression values, STimage also implements a classification model that enables the prediction of high vs low gene expression directly from H&E images (Figure S11c). Refer to Section 2.6 for details on the classification model. In this model, continuous gene expression values were discretised into two categories: high vs low (Figure S11a,b).

We evaluated the classification model’s performance on the top 100 predictable genes in two test samples—FFPE and 1160920F—using Moran’s I, AUC, and IoU metrics (Figure S11c). As the aim of this model is to classify tissue regions into high vs low expression zones, IoU is the most informative metric. The top predictable genes showed consistent classification performance across both test samples (Figure S11d,e). Since the classification model predicts at the tile level, we found that it performed well on tiles with high expression, but was noisier for tiles with intermediate expression levels. Examples of top predictable genes include CD24, GNAS, and VEGFA (Figure S11d,e). These genes are highly relevant to cancer; for instance, CD24 is known to be overexpressed in HER2+ tumours and is associated with poor prognosis^43^. To assess the classification model’s robustness, we compared two genes with contrasting expression distributions—CD52 and CD24 (Figure S11). CD52 is an immune-related gene and a favourable biomarker in breast cancer^44^. Class probabilities from the model were consistent with the ground truth continuous gene expression values (Figure S11g). All three performance metrics supported the model’s predictive accuracy for both genes.

### 3.11 Interpretability analysis

We investigated the interpretability of the model by assessing feature importance using Linear Interpretability Model-agnostic Explanations (LIME) (Figure S12; see Methods, Section 2.5). As examples, we selected VEGFA and CD52, as they were ranked among the top predictable genes and are known cancer and immune markers, respectively. We calculated LIME scores for the regression models (FFPE and 1160920F), focusing on the top five tiles with high VEGFA expression from cancer regions and the top five tiles with high CD52 expression from immune regions (Figure S12c,d). LIME scores overlaid on H&E images, used for both qualitative and quantitative assessment, are shown in Figure S12a,b. Two representative examples are displayed for two different test datasets: FFPE (Figure S12a) and 1160920F (Figure S12b).

The interpretability analysis shows that different nuclei contribute differently to gene expression prediction (Figure S12a,b). We observed that nuclei with positive LIME scores for VEGFA have a larger area than those with positive LIME scores for CD52 in cancer regions, consistent with the biological expectation that cancer cells are typically larger than immune cells. When integrated with pathological annotations, we also found more nuclei with high LIME scores for VEGFA in cancer tiles (Figure S12a,b) than in immune tiles. This observation is consistent with the known biology that VEGFA is overexpressed in cancer cells in HER2+ samples^45^. To support user engagement and exploration of interpretability, we developed an interactive web application that allows visualisation of LIME scores from the STimage model on arbitrary tiles.

Moreover, analysis of the latent space revealed that the learned features from ResNet50 could effectively distinguish cancer tiles from non-cancer tiles. We overlaid the true expression and predicted probability of CD24—one of the top predictable genes across both datasets—on the first two principal components of the latent space (Figure S11f). CD24 is highly expressed in cancer regions and aligns well with pathologist annotations in Figure S12a,b. In addition, saliency mapping confirmed that ResNet50 focuses on the most informative features (nuclei) in the images (Figure S12).

### 3.12 Xenium cell type classification

The Xenium platform from 10x Genomics enables RNA transcript detection with subcellular resolution. Compared to Visium, Xenium-generated data is particularly well-suited for deep learning-based cell classification due to its single-cell resolution profiling. We obtained Xenium data for five primary breast cancer tissue sections and trained a deep learning model to classify nuclei using whole-slide H&E images.

Figure 4 shows the classification metrics (Figure 4a,b), indicating that cancer epithelial cells were the most accurately predicted, while other cell types also showed reasonable accuracy. Qualitatively, predictions closely matched the ground truth labels at both the macroscopic scale (Figure 4d), where distinct tissue structures were well reconstituted, and the microscopic scale (Figure 4e), showing individual nuclei segmentation and classification.

**Figure 4.**
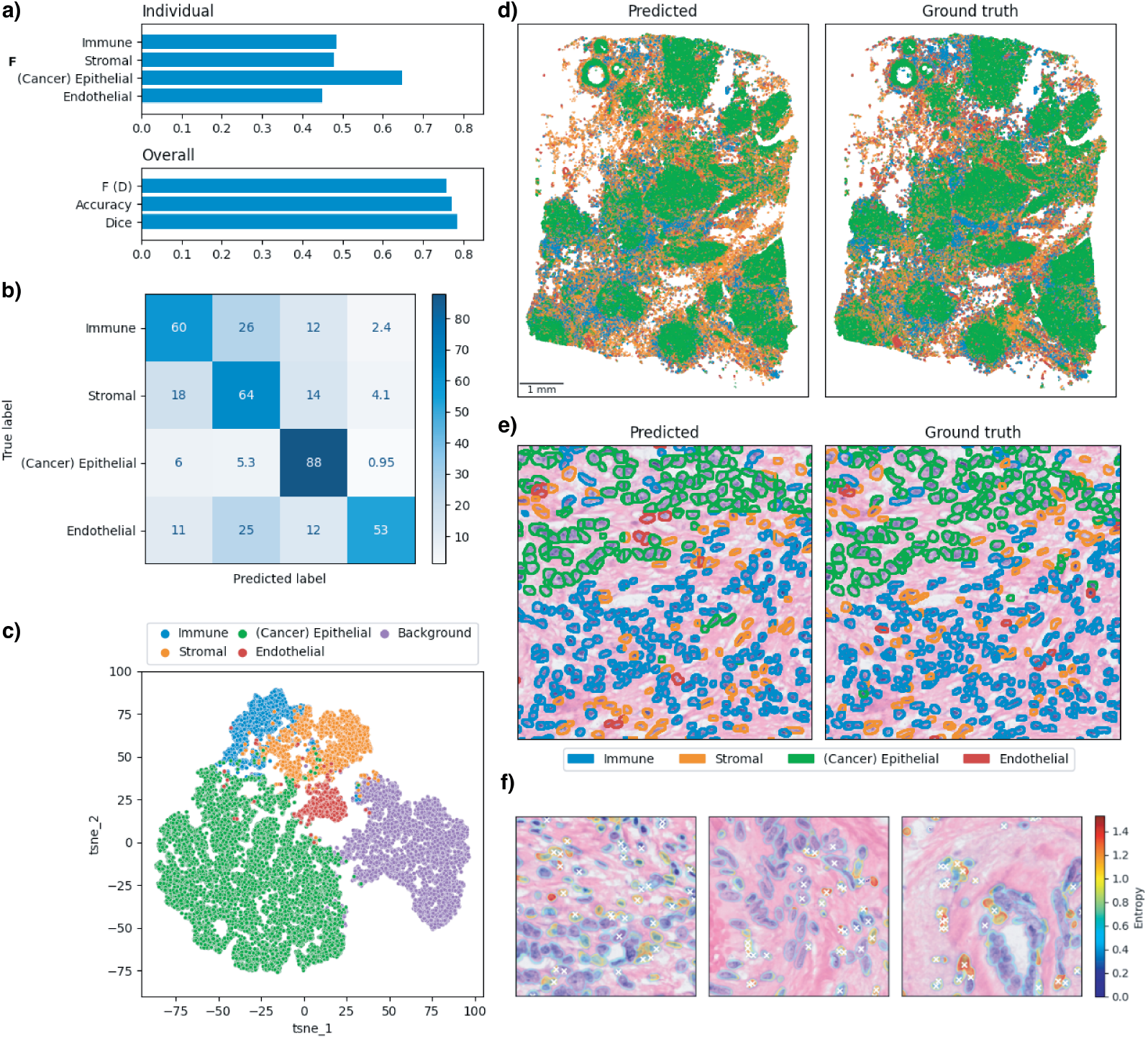
Cell type classification at single-cell resolution using a customised Hover-Net model on Xenium data. **a**, Classification and segmentation metrics evaluated on the held-out test sample. The top panel shows individual classification F1 scores for each cell type. The bottom panel reports overall metrics: F (D) is the detection F1 score, accuracy refers to the classification accuracy within correctly detected instances across all cell types, and the Dice coefficient measures segmentation quality. **b**, Confusion matrix for correctly detected instances, showing percentages normalised across the true labels (rows). **c**, Visualisation of the features corresponding to predicted cell types using t-SNE. **d**, Predicted and ground truth cell types across the full held-out test whole-slide image. **e**, Close-up example of segmentation and classification of nuclei in the test image, comparing predictions to ground truth. **f**, Visualisation of prediction uncertainty, measured by the entropy of predicted cell class probabilities. White crosses indicate misclassified instances.

We also visualised the feature space learned by the model to interpret inter-cell type similarity and dissimilarity. The 256 × 256 × 64 feature map from the final layer of the classification branch was associated with the predicted cell types. Feature vectors were sampled for each cell type and the background class, and visualised using 2D t-SNE embeddings (Figure 4c). The background was clearly separated from cells, while stromal, immune, and endothelial cells showed substantial overlap, suggesting potential classification challenges among these types.

Furthermore, we assessed model uncertainty by computing the entropy of predicted class probabilities across the image. We overlaid this uncertainty map with misclassified instances. Figure 4f shows that regions of high entropy were enriched for misclassified cells, while low-entropy regions largely corresponded to correct predictions.

We performed leave-one-out cross-validation (LOOCV) across the five Xenium datasets. Results presented in Figure S14 show consistent performance across test samples, demonstrating the robustness of our classification model. Among the cell types, cancer epithelial cells were most accurately predicted, likely due to their abundance and distinctive morphology. Immune cells (including B cells, T cells, and myeloid cells) formed the second most predictable group. This suggests potential clinical applications in profiling immune cell distribution in the tumour microenvironment, which is critical for cancer diagnosis. With more training data, particularly for rarer cell types, predictive performance could be further improved.

We benchmarked our model against the recently published CellViT model^46^, a Vision Transformer-based architecture that supports both single-cell segmentation and classification (Figure S15). At the tile level, both CellViT and the STimage classification model accurately predicted cell segments. However, the STimage model outperformed CellViT in cell type prediction (Figure S15a). We also compared mean IoU scores across LOOCV test samples (Figure S15b), demonstrating that our model achieved superior performance.

Notably, the STimage classification model was trained using Xenium data in which training labels were generated automatically via image registration. This process aligns the DAPI-derived cell segmentation mask with cell type annotations obtained from marker gene expression measurements in the Xenium dataset, which are considered more accurate than those inferred from H&E alone. This automated approach reduces the need for manual labelling and minimises human error.

### 3.13 Evaluating clinical and translational potential of STimage using the TCGA dataset and a drug response dataset

To evaluate whether STimage could be applied to tissue images from a non-spatial dataset, we first tested two randomly selected breast cancer samples from The Cancer Genome Atlas (TCGA) (Figure 1f,g,h). As ground truth for comparison, we obtained pathological annotations for these two H&E images. The predictions for cancer markers were consistent across the two TCGA replicates (Figure 1f,g,h). We observed strong agreement between the predicted expression and pathological annotation, with high expression of cancer markers in tumour regions and low expression outside tumour areas.

We further demonstrate that the model-predicted gene expression can cluster cell types from H&E images, even in the absence of spatial measurements. This was evaluated using five tissue sections from three randomly selected TCGA patients. Louvain clustering based on the expression of 1,522 predicted genes per tile is shown in Figure S13a. The resulting clusters correlated with pathological annotations for each sample (Figure S13a,b). Spatial plots of two known breast cancer markers—ESR1 and GATA3^47^—also aligned with annotated tumour and normal regions (Figure S13c,d).

For a broader analysis, we predicted expression levels of 1,522 highly variable genes for 1,034 H&E images from 670 TCGA patients. The average Pearson correlation coefficient between predicted and bulk-measured gene expression was 0.48 (Figure 5a). STimage performed consistently well for both lowly expressed genes (e.g., MPPED1) and highly expressed genes (e.g., VEGFA), as illustrated in Figure 5b. Notably, the predicted gene expression was also informative for patient survival (see Methods, Section 2.10). Using the top five genes identified from univariate Cox regression models, we predicted gene expression for 1,034 images with matched survival data. Survival curves stratifying patients into low- and high-risk groups were significant and consistent for both true and predicted expression values (Figure 5c,d).

**Figure 5.**
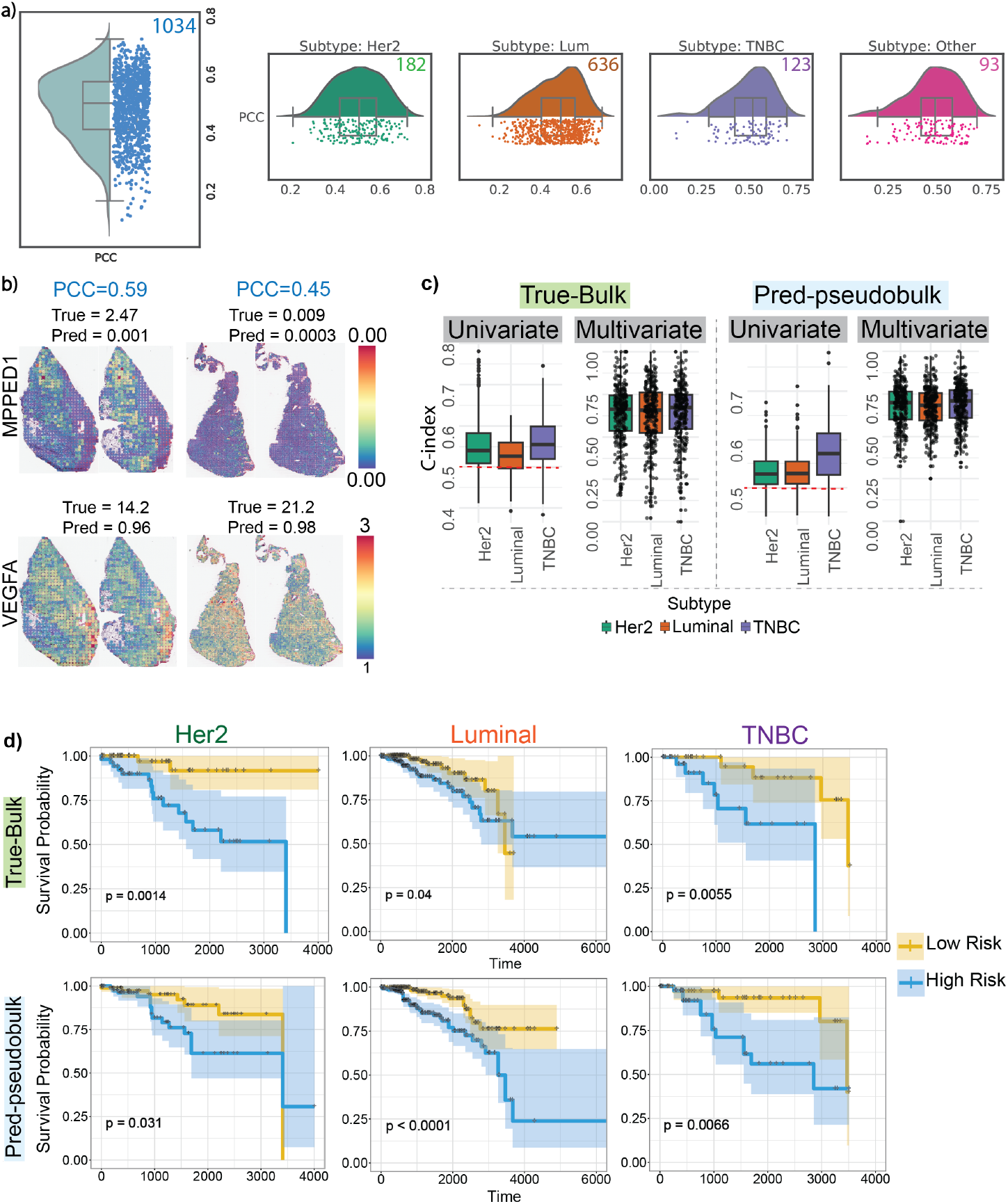
TCGA validation. **a,** Box plot showing gene-wise PCC per sample, computed between predicted gene expression from TCGA H&E images and matched bulk gene expression across 1,034 samples/images from 670 patients. Each dot represents a patient. The accompanying violin plots correspond to individual subtypes; the number in the top right of each indicates the number of samples in that subtype. **b,** Spatial gene expression from the Visium dataset for a highly expressed gene (VEGFA) and a lowly expressed gene (MPPED1) for two randomly selected patients. **c,** Box plots showing performance of prognostic models using the c-index for true and predicted data across univariate and multivariate models. **d,** Kaplan–Meier survival curves stratifying patients into low- and high-risk groups, based on true (measured bulk RNA-seq) and predicted gene expression from images for TCGA samples. Multivariate models based on both true and predicted data captured significant differences between low- and high-risk groups.

Top genes with the highest PCCs from gene-wise correlation analysis for each subtype were: FGFR3 (PCC = 0.46) and PAK1 (PCC = 0.47) for HER2; IL5RA (PCC = 0.19) and CCL13 (PCC = 0.20) for Luminal; and KLF4 (PCC = 0.47) and BDNF (PCC = 0.48) for TNBC. These genes are known to be associated with treatment resistance or breast cancer progression. For example, PAK1 copy number is increased in HER2-positive cases^48^, and its overexpression is linked to poor prognosis and metastasis^49^. KLF4 is a known prognostic marker in TNBC^50^.

To further evaluate the clinical utility of STimage, we independently assessed its ability to predict responses to drugs. Here, we analysed another cohort of breast cancer patients with treatment response information^51^. Average gene-wise Pearson’s correlation coefficient was found to be 0.47 for 1139 common genes (Figure S16b). We then evaluated the performance of true and predicted gene expression of 1139 common genes for classifying pathological complete response (pCR) vs residual disease (RD) using an RBF-kernel support-vector machine (SVM) classifier. Given the different distribution of the data, the predicted gene expression data was log-transformed, while the true raw gene expression data was CPM-normalised and standardized.

We used a two-level stratified cross-validation approach to test how well our model could predict new data. First, we divided the full dataset into five equal folds. Each time, we used four folds to train the model and the remaining fold as held-out set. Within the four folds, we did another round of stratified 5-fold splitting, to get the best trained model to be used for predicting the 5th held out set. We repeated this process five times so that every patient was part of the held-out set. This was done to ensure the results are robust and reliable. STimage could stratify patient responses with AUC = 0.70, compared with using the bulk RNA-seq with AUC = 0.78 (Figure S16c).

## 4 Discussions and Future perspectives

While digital pathology has the potential to benefit greatly from spatial tissue analysis, it still faces significant challenges related to cost, time efficiency, reliability, and explainability^52^. We developed STimage, a comprehensive machine learning software to predict spatial gene expression and cell types accurately by training on a small set of spatial transcriptomics data, and subsequently applying the model to standard H&E images. This cost- and time-efficient approach is able to predict genes and cell types biologically relevant to the disease, while maintaining reliability across datasets and interpretability.

The STimage regression model integrates deep learning and statistical approaches to estimate the distribution of gene expression in spatial transcriptomics (ST) data and quantify prediction uncertainty. We benchmarked the STimage regression model against existing methods in both leave-one-out cross-validation experiments and out-of-distribution settings. In most cases, STimage outperformed other methods. Additionally, STimage includes a classification model that can predict high or low expression levels for a gene or gene list. STimage also implements a LIME-based interpretability model to identify meaningful tissue and cellular features that contribute to model predictions.

As spatial data remains limited due to high costs and lack of access to the spatial platforms, the datasets used in this study are modest in size. Nonetheless, we used highly information-rich data, comprising 87,166 spatial spots and 2.1 million cells from 76 patients, in addition to 670 patients from the TCGA. Despite the modest sample size, the study comprehensively covers different spatial resolutions (from single-cell to large-spot scale), protocols (polyA-capture, probe-based, and imaging-based), sample types (fresh frozen and FFPE), molecular modalities (RNA and protein), cancer types (skin, breast, and kidney cancer), and non-spatial data. This diversity enabled a robust assessment of model generalisability.

Although STimage outperformed existing tools, Pearson correlation values remain modest overall. However, spatial expression patterns are arguably more biologically relevant. Therefore, we propose a new evaluation metric for spatial distribution, Intersection over Union (IoU), to better reflect spatial prediction accuracy. This metric suggests impressive performance for STimage and potential utilities in clinical settings. Notably, STimage identifies the most predictable genes, which are functionally relevant to the respective cancer types, demonstrated as relevant enriched pathways for skin cancer, lung cancer, kidney cancer, breast cancer, as well as non-cancer liver disease. For example, SOX10 (a highly sensitive and specific nuclear marker for melanocytic lesions, including primary and metastatic melanoma, PRAME (overexpressed in many melanomas but not in benign nevi or normal melanocytes, a marker and a target for immunotherapy and vaccine-based approaches in melanoma treatment), and S100B (a sensitive marker for melanocytic differentiation in tissue samples) were the top genes with the highest prediction accuracy for melanoma. Similarly, for breast cancer, the top predictable genes include ESR1 (major driver in hormone receptor positive breast cancer), HER2 (ERBB2, a member of the EGFR family of receptor tyrosine kinases, associated with higher cancer progression and recurrence), GATA3 (strong correlation with ER-positive tumours and favourable prognosis), and immune markers like CD45 (marking immune infiltration). We assessed potential clinical utilities through survival analyses and patient stratification for drug responses. The predicted results based on the images were able to classify long vs short survival and complete vs partial responses. While interpretability measures are still in early development, initial results suggest informative latent space embeddings that can differentiate pathological annotations and identify predictive nuclear features.

Overall, we believe STimage represents meaningful progress in applying machine learning to digital pathology and provides a practical, interpretable, and extensible tool for spatial transcriptomics-based gene expression and cell type prediction.

## 5 Declarations

### 5.1 Ethics approval and consent to participate

Not applicable.

### 5.2 Availability of data and materials

The datasets analysed in this study are publicly available and can be downloaded from UQ eSpace via https://doi.org/10.48610/4fb74a9. The STimage software and web-app tools are available on GitHub at https://github.com/BiomedicalMachineLearning/STimage.

## 5.3 Acknowledgements

We thank Prof. Alex Swarbrick for kindly providing the spatial transcriptomics data. We also thank the staff at the University of Queensland Sequencing Facility and the School of Biomedical Sciences Imaging Facility for their support with spatial transcriptomics sequencing.

## 5.4 Funding

This research was partially supported by the Australian Research Council through the Industrial Transformation Training Centre for Information Resilience (IC200100022). Q.H.N. is supported by the Australian Research Council (ARC DECRA Grant DE190100116), the National Health and Medical Research Council (NHMRC Project Grant 2001514), and the NHMRC Investigator Grant (GNT2008928).

### 5.6 Author contributions

Q.H.N. and X.T. conceived the project. Q.H.N., X.T., O.M., and S.M. designed the algorithm. S.M. led the uncertainty quantification analysis. X.T. and O.M. developed the software. K.T. and P.S. performed pathological annotations. F.R., M.T., and N.Y. contributed to the data analysis. All authors contributed to the writing of the manuscript.

## 6 Supplementary data

## References

1. Elmore, J. G. et al. Pathologists’ diagnosis of invasive melanoma and melanocytic proliferations: observer accuracy and reproducibility study. BMJ 357, j2813 (2017).

2. Patel, A. P. et al. Single-cell RNA-seq highlights intratumoral heterogeneity in primary glioblastoma. Science 344, 1396–1401 (2014).

3. Evans, A. J. et al. US food and drug administration approval of whole slide imaging for primary diagnosis: A key milestone is reached and new questions are raised. Arch. Pathol. Lab. Med. 142, 1383–1387 (2018).

4. Saltz, J. et al. Spatial organization and molecular correlation of tumor-infiltrating lymphocytes using deep learning on pathology images. Cell Rep. 23, 181–193.e7 (2018).

5. Esteva, A. et al. Dermatologist-level classification of skin cancer with deep neural networks. Nature 542, 115–118 (2017).

6. Janda, M. & Soyer, H. P. Can clinical decision making be enhanced by artificial intelligence? Br. J. Dermatol. 180, 247–248 (2019).

7. Schmauch, B. et al. A deep learning model to predict RNA-Seq expression of tumours from whole slide images. Nat. Commun. 11, 3877 (2020).

8. Ståhl, P. L. et al. Visualization and analysis of gene expression in tissue sections by spatial transcriptomics. Science 353, 78–82 (2016).

9. He, B. et al. Integrating spatial gene expression and breast tumour morphology via deep learning. Nat. biomedical engineering 4, 827–834 (2020).

10. Pang, M., Su, K. & Li, M. Leveraging information in spatial transcriptomics to predict super-resolution gene expression from histology images in tumors. bioRxiv DOI: 10.1101/2021.11.28.470212 (2021). https://www.biorxiv.org/content/early/2021/11/28/2021.11.28.470212.full.pdf.

11. Zeng, Y. et al. Spatial transcriptomics prediction from histology jointly through transformer and graph neural networks. Brief. Bioinform. 23 (2022).

12. Monjo, T., Koido, M., Nagasawa, S., Suzuki, Y. & Kamatani, Y. Efficient prediction of a spatial transcriptomics profile better characterizes breast cancer tissue sections without costly experimentation. Sci. Rep. 12, 4133 (2022).

13. Doshi-Velez, F. & Kim, B. Towards a rigorous science of interpretable machine learning (2017). 1702.08608.

14. Wu, S. Z. et al. A single-cell and spatially resolved atlas of human breast cancers. Nat. Genet. 53, 1334–1347 (2021).

15. Raghubar, A. M. et al. High risk clear cell renal cell carcinoma microenvironments contain protumour immunophenotypes lacking specific immune checkpoints. npj Precis. Oncol. 7, 88, DOI: 10.1038/s41698-023-00208-9 (2023).

16. Andrews, T. S. et al. Single-cell and spatial transcriptomics characterisation of the immunological landscape in the healthy and psc human liver. J. Hepatol. DOI: 10.1016/j.jhep.2023.12.020 (2024).

17. Vahadane, A. et al. Structure-preserving color normalization and sparse stain separation for histological images. IEEE Trans. Med. Imaging 35, 1962–1971 (2016).

18. He, K., Zhang, X., Ren, S. & Sun, J. Deep residual learning for image recognition, DOI: 10.48550/ARXIV.1512.03385 (2015).

19. Ribeiro, M. T., Singh, S. & Guestrin, C. “why should I trust you?”: Explaining the predictions of any classifier. CoRR abs/1602.04938 (2016). 1602.04938.

20. Stringer, C., Wang, T., Michaelos, M. & Pachitariu, M. Cellpose: a generalist algorithm for cellular segmentation. Nat. methods 18, 100–106 (2021).

21. Zou, H. & Hastie, T. Regularization and variable selection via the elastic net. J. royal statistical society: series B (statistical methodology) 67, 301–320 (2005).

22. Defazio, A., Bach, F. & Lacoste-Julien, S. Saga: A fast incremental gradient method with support for non-strongly convex composite objectives. Adv. neural information processing systems 27 (2014).

23. Graham, S. et al. Hover-net: Simultaneous segmentation and classification of nuclei in multi-tissue histology images. Med. image analysis 58, 101563 (2019).

24. Lowe, D. G. Distinctive image features from scale-invariant keypoints. Int. journal computer vision 60, 91–110 (2004).

25. Hörst, F. et al. CellViT: Vision Transformers for precise cell segmentation and classification. Med. Image Analysis 94, 103143, DOI: 10.1016/j.media.2024.103143 (2024). Epub 2024 Mar 16.

26. Clark, K. et al. The cancer imaging archive (tcia): maintaining and operating a public information repository. J. digital imaging 26, 1045–1057 (2013).

27. Xu, J. et al. Delving into the heterogeneity of different breast cancer subtypes and the prognostic models utilizing scrna-seq and bulk rna-seq. Int. J. Mol. Sci. 23, 9936 (2022).

28. Therneau, T. M. A Package for Survival Analysis in R (2024). R package version 3.5-8.

29. Strbenac, D., Mann, G. J., Ormerod, J. T. & Yang, J. Y. Classifyr: an r package for performance assessment of classification with applications to transcriptomics. Bioinformatics 31, 1851–1853 (2015).

30. Sammut, S.-J. et al. Multi-omic machine learning predictor of breast cancer therapy response. Nature 601, 623–629, DOI: 10.1038/s41586-021-04278-5 (2022).

31. Bergenstråhle, L. et al. Super-resolved spatial transcriptomics by deep data fusion. Nat. Biotechnol. 40, 476–479 (2022).

32. Andersson, A. et al. Spatial deconvolution of her2-positive breast cancer delineates tumor-associated cell type interactions. Nat. communications 12, 6012 (2021).

33. Xu, H., Usuyama, N., Bagga, J. et al. A whole-slide foundation model for digital pathology from real-world data. Nature 630, 181–188, DOI: 10.1038/s41586-024-07441-w (2024).

34. Bioptimus. Bioptimus launches h-optimus 0: the world’s largest open-source ai foundation model for pathology. Bioptimus News (2024).

35. Patel, A., Xu, X., Van, T. N. & Fuchs, T. J. Virchow2: Scaling multi-resolution vision transformers for histopathology. arXiv preprint 2408.00738 (2024).

36. Filiot, A., Jacob, P., Mac Kain, A. & Saillard, C. Phikon-v2: A large and public feature extractor for biomarker prediction. arXiv preprint 2409.09173 (2024).

37. Wang, X. et al. Transformer-based unsupervised contrastive learning for histopathological image classification. Med. Image Analysis 81, 102559, DOI: 10.1016/j.media.2022.102559 (2022).

38. Lu, M. et al. A visual-language foundation model for computational pathology. Nat Med 30, 863–874, DOI: 10.1038/s41591-024-02856-4 (2024). Epub 2024 Mar 19.

39. Chen, R. J., Ding, T., Lu, M. Y. et al. Towards a general-purpose foundation model for computational pathology. Nat Med 30, 850–862, DOI: 10.1038/s41591-024-02857-3 (2024).

40. Vorontsov, E., Bozkurt, A., Casson, A. et al. A foundation model for clinical-grade computational pathology and rare cancers detection. Nat. Medicine 30, 2924–2935, DOI: 10.1038/s41591-024-03141-0 (2024).

41. MacDonald, S., Steven, K. & Trzaskowski, M. Interpretable ai in healthcare: Enhancing fairness, safety, and trust. In Artificial Intelligence in Medicine: Applications, Limitations and Future Directions, 241–258 (Springer, 2022).

42. MacDonald, S. et al. Generalising uncertainty improves accuracy and safety of deep learning analytics applied to oncology. bioRxiv 2022–07 (2022).

43. Hosonaga, M., Arima, Y., Sugihara, E., Kohno, N. & Saya, H. Expression of cd 24 is associated with her2 expression and supports her2-akt signaling in her2-positive breast cancer cells. Cancer science 105, 779–787 (2014).

44. Ma, Y.-F. et al. The immune-related gene cd52 is a favorable biomarker for breast cancer prognosis. Gland Surg. 10, 780 (2021).

45. Al Kawas, H. et al. How vegf-a and its splice variants affect breast cancer development–clinical implications. Cell. Oncol. 45, 227–239 (2022).

46. Hörst, F. et al. Cellvit: Vision transformers for precise cell segmentation and classification (2023). 2306.15350.

47. McCleskey, B. C. et al. Gata3 expression in advanced breast cancer: prognostic value and organ-specific relapse. Am. journal clinical pathology 144, 756–763 (2015).

48. Skjervold, A. H., Valla, M., Ytterhus, B. & Bofin, A. M. Pak1 copy number in breast cancer—associations with proliferation and molecular subtypes. PLoS One 18, e0287608 (2023).

49. Belli, S. et al. Pak1 pathway hyper-activation mediates resistance to endocrine therapy and cdk4/6 inhibitors in er+ breast cancer. NPJ Breast Cancer 9, 48 (2023).

50. Nagata, T. et al. Klf4 and nanog are prognostic biomarkers for triple-negative breast cancer. Breast cancer 24, 326–335 (2017).

51. Sammut, S.-J. et al. Multi-omic machine learning predictor of breast cancer therapy response. Nature 601, 623–629 (2022).

52. Pang, J.-M. B. et al. Spatial transcriptomics and the anatomical pathologist: Molecular meets morphology. Histopathology 84, 577–586, DOI: 10.1111/his.15093 (2024). https://onlinelibrary.wiley.com/doi/pdf/10.1111/his.15093.

53. Xu, C. et al. S100a14, a member of the ef-hand calcium-binding proteins, is overexpressed in breast cancer and acts as a modulator of her2 signaling. J. Biol. Chem. 289, 827–837 (2014).

