## supplemental text and figure for "Robust and interpretable prediction of gene markers and cell types from spatial transcriptomics data"

| No. | Dataset | Samples | Tissues | Resolution | Unfiltered spots | Remaining spots | Genes |
|---|---|---|---|---|---|---|---|
| 1 | Her2ST[32] | 36 | Frozen | 100 $\mu M$ | 13,653 | 12,584 | 11,871 |
| 2 | Swarbrick's Lab[14] | 6 | Frozen | 55 $\mu M$ | 15,611 | 14,951 | 14,664 |
| 3 | Public BC 10X-Visium | 2 | Frozen | 55 $\mu M$ | 7,785 | 7,289 | 36,601 |
| 4 | Public BC 10X-Visium | 1 | FFPE | 55 $\mu M$ | 2,518 | 2,338 | 36,601 |
| 5 | Melanoma | 13 (5 slides) | FFPE | 55 $\mu M$ | 15,209 | 14,405 | 17,943 |
| 6 | Liver Visium[16] | 4 | Frozen | 55 $\mu M$ | 19,968 | 19,967 | 36,601 |
| 7 | KC Visium[15] | 6 | Frozen | 55 $\mu M$ | 16,131 | 15,632 | 36,601 |
| 8 | BC Xenium | 5 | Frozen | subcellular | - | 21,076 (1,294,600 cells) | 280 |
| 9 | SC CODEX | 3 (1 slide) | Frozen | subcellular | - | 6,174 (813,004 cells) | 34 (Proteins) |

**Table 1.** Overview of Datasets used by STimage.
BC - Breast Cancer, KC - Kidney Cancer, SC - Skin Cancer

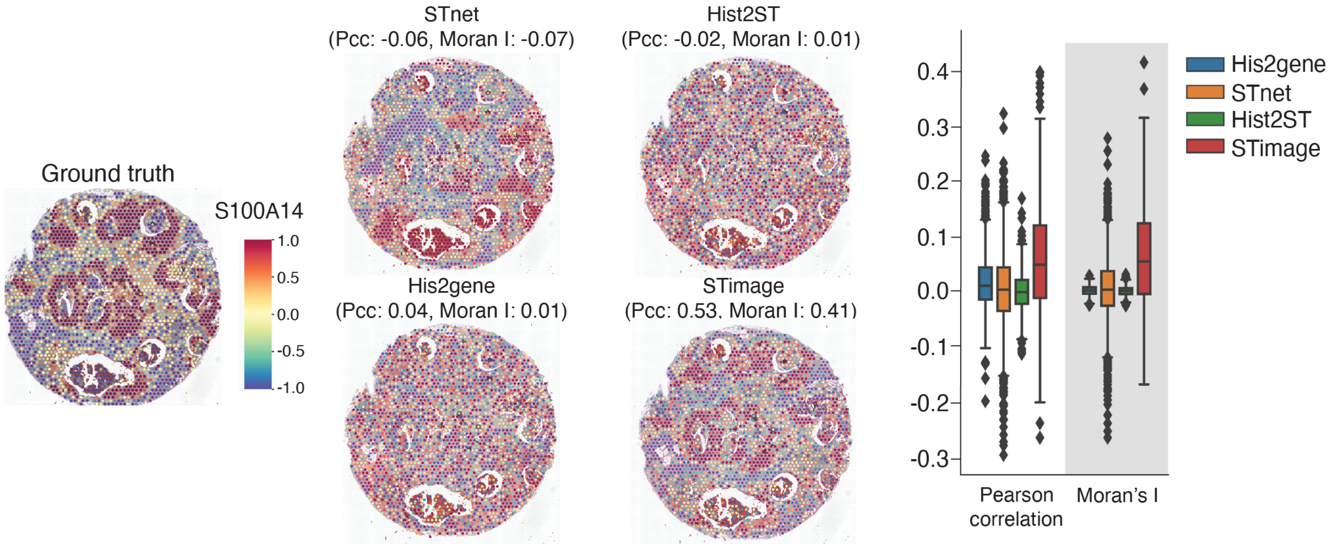


**Figure S1.** Benchmarking of the STimage regression model against STnet, HisToGene, and Hist2ST models using Visium data from 10x Genomics. All models were trained on two fresh frozen breast cancer tissue samples and tested on an out-of-distribution (OOD) FFPE tissue. The models were intentionally trained and evaluated using a small dataset and OOD samples to assess robustness. Visualisation compares the ground truth expression of the cancer marker S100A14[53] with predicted expression from the different models. PCC indicates the Pearson correlation coefficient, and MI refers to Moran's I (spatial autocorrelation). PCC and Moran's I scores for the predicted 1,000 highly variable genes (HVGs) were compared against ground truth for each model.

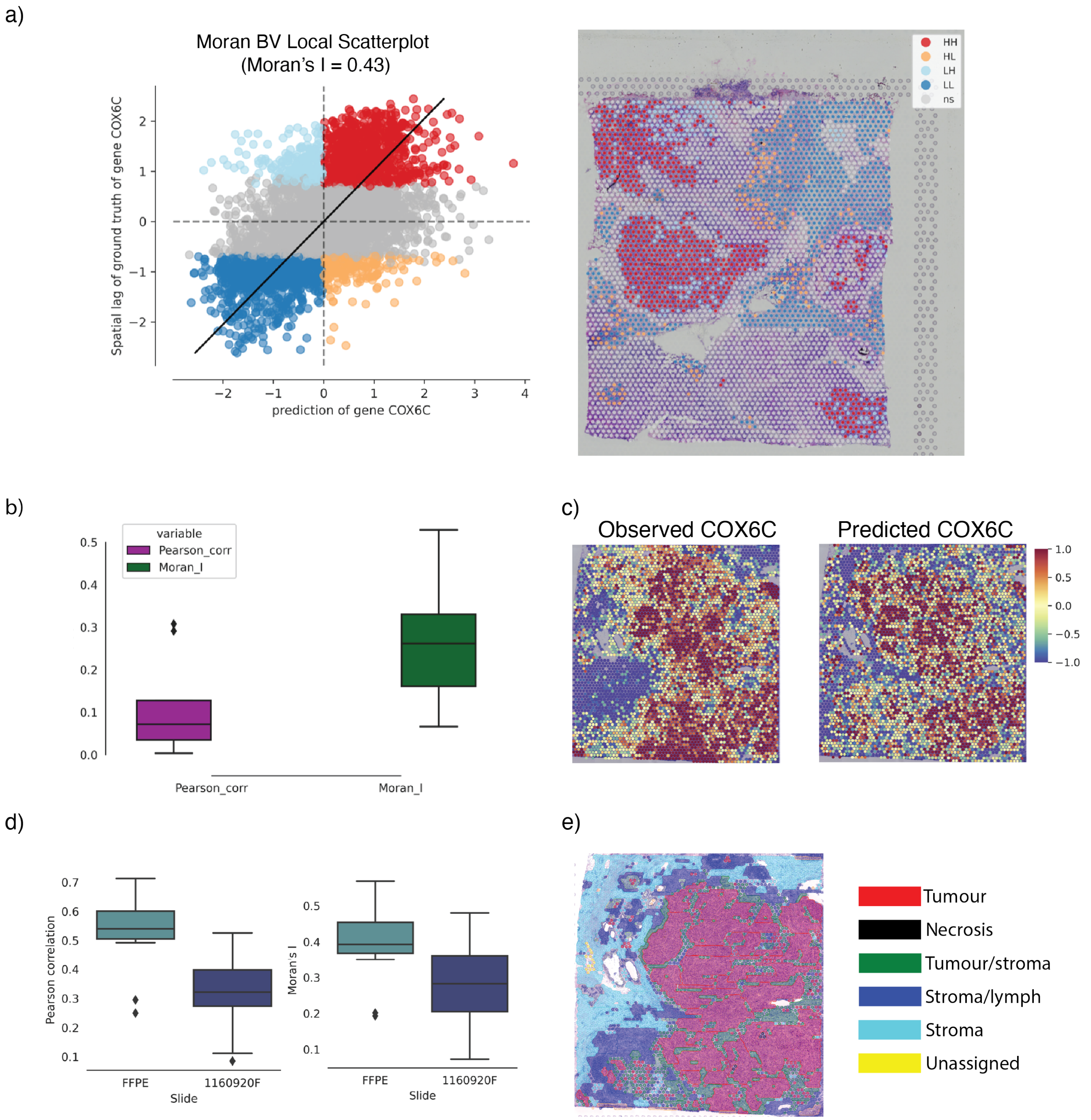


**Figure S2.** Performance of the STimage CNN–NB regression model. **a**, Spatial autocorrelation using Moran's I index for the predicted expression of the highly abundant cancer marker COX6C. The points in the scatter plot correspond to those shown in the tissue plot. HH indicates concordance between high predicted and high observed values; LL = Low–Low, HL = High–Low, LH = Low–High, and ns = non-significant. **b**, PCC and Moran's I scores calculated for predicted and observed expression of 14 cancer and immune markers. **c**, Model performance on an external test dataset. **d**, PCC and Moran's I values for predicted and observed expression of 14 cancer and immune markers in the external test dataset. **e**, Ground truth cell type labels for the external test dataset, defined by a pathologist as described in[14].

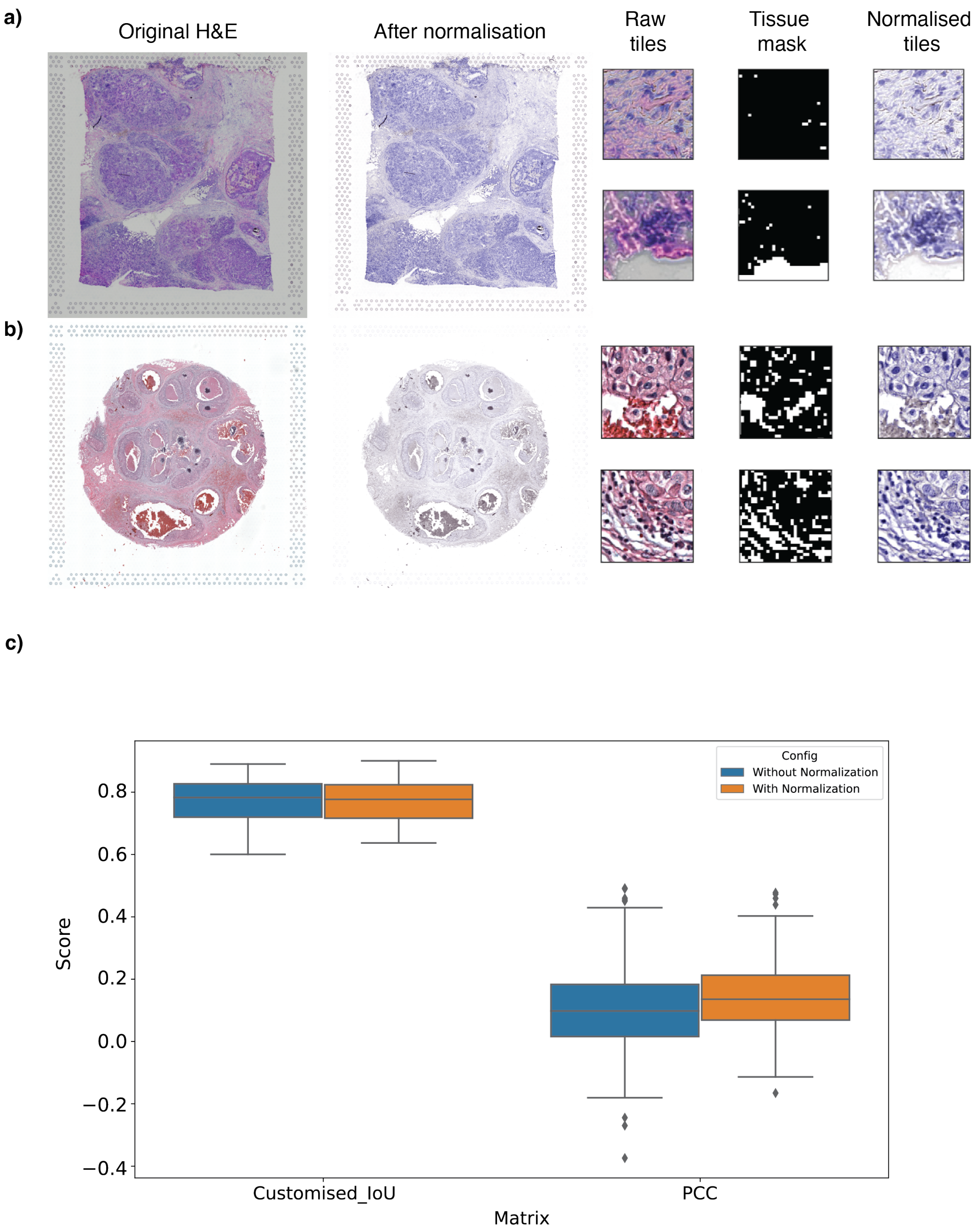


**Figure S3.** Preprocessing of H&E image input prior to running the STimage model. **a**, Normalisation and tile removal for fresh frozen tissue samples. **b**, As in (a), but for FFPE tissue. Tissue masking was performed using OpenCV2, where black areas in the image indicate detected tissue, and white areas represent regions without tissue coverage. Tiles with less than 70% tissue coverage were removed. Vahadane stain normalisation[17] was applied (as the default method) to correct for background staining across all images to match a template image. Images before and after normalisation are shown. **c**, Comparison of model performance with and without stain normalisation, shown as a PCC evaluation matrix.

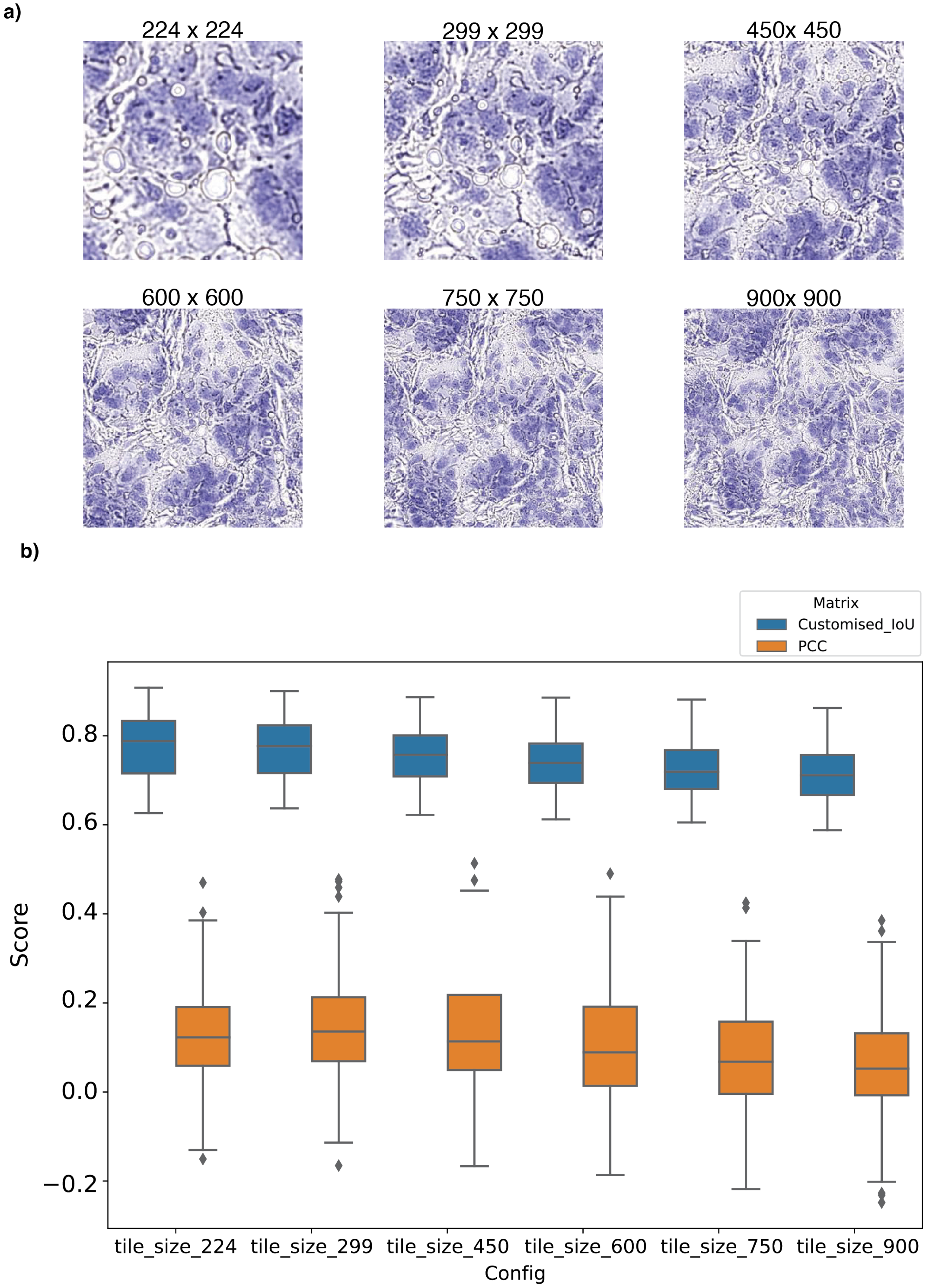


**Figure S4.** Optimisation of tile size and its effect on model performance based on the breast cancer Visium dataset. **a**, Tile size options included 244 × 244, 299 × 299, 450 × 450, 600 × 600, 750 × 750, and 900 × 900. **b**, Model performance across different tile sizes was evaluated using Pearson correlation coefficient (PCC) and the geometry-based Intersection over Union (IoU) metric.

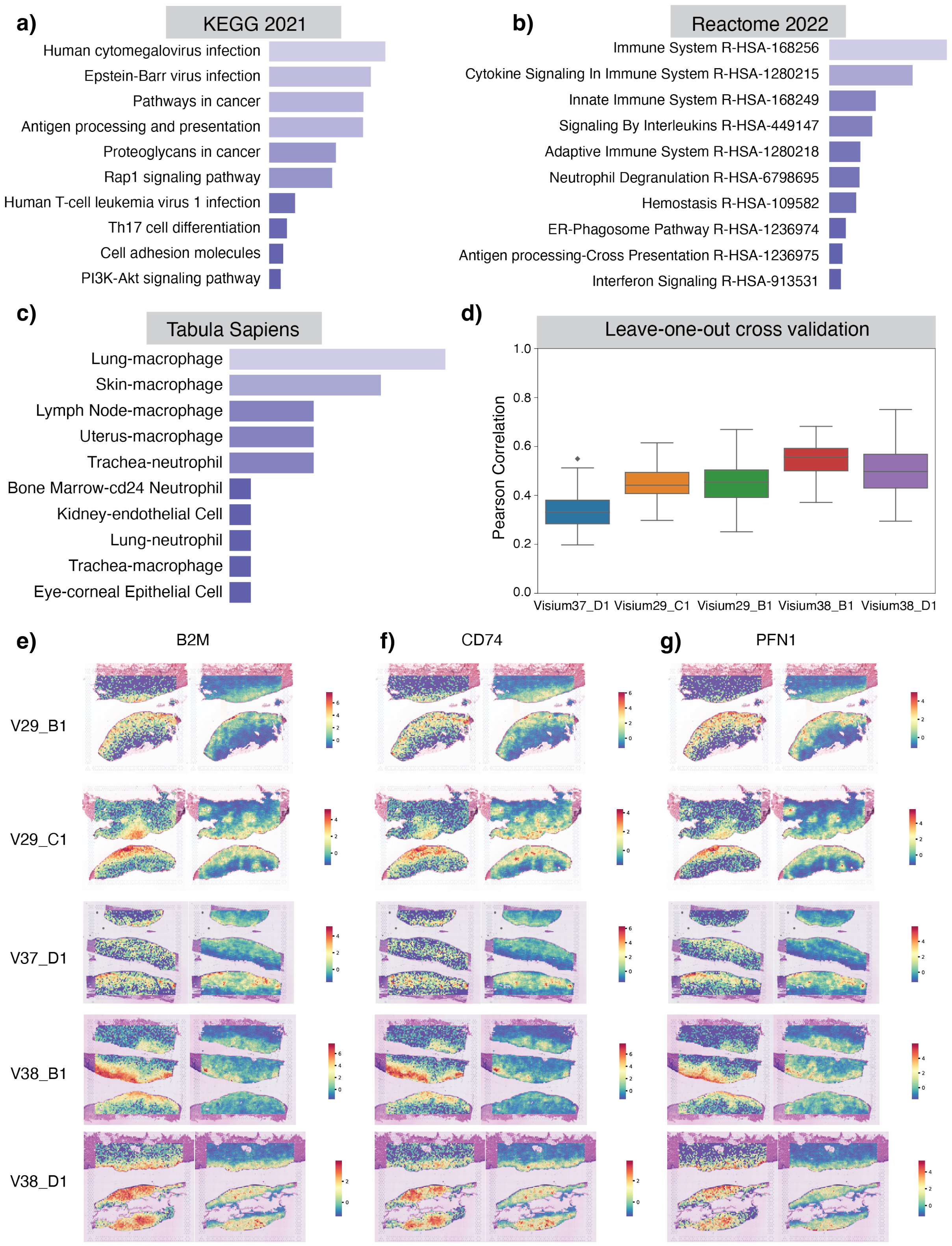


**Figure S5.** Performance of the STimage regression model on a skin cancer dataset in a Leave-One-Out Cross-Validation (LOOCV) experiment. **a, b, c**, Top GO enrichment terms from the KEGG 2021, Reactome 2022, and Tabula Sapiens databases, based on the top 100 most predictable genes identified by the STimage model. **d**, PCC between STimage-predicted gene expression and ground truth measurements from the Visium experiment in a LOOCV setting, shown for each test sample. **e, f, g**, Tissue plots showing ground truth gene expression for three skin cancer-related genes from the top 100 most predictable genes. Each row represents a leave-one-out test sample.

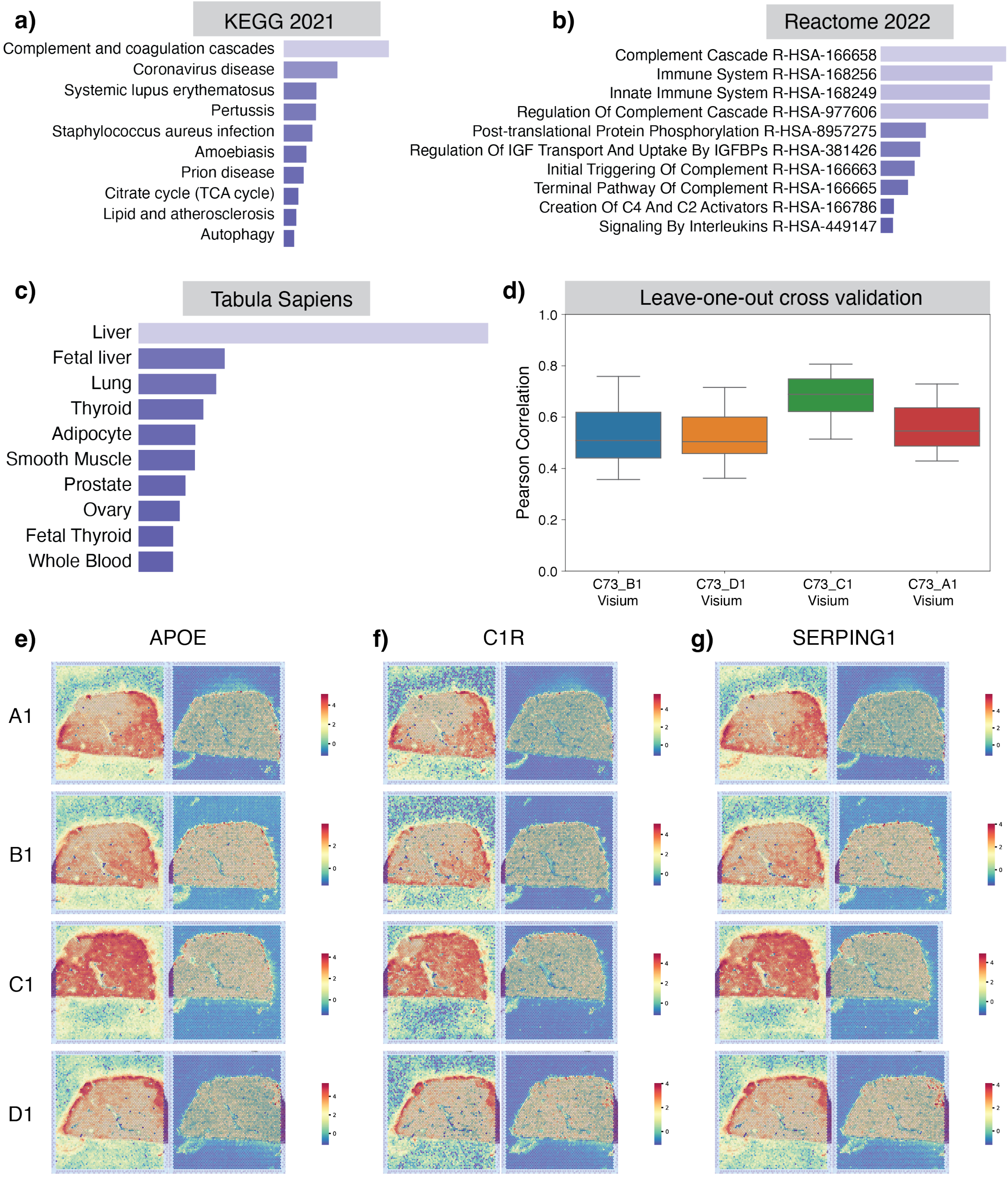


**Figure S6.** Performance of the STimage regression model on a liver dataset in a LOOCV experiment. **a, b, c**, Top GO enrichment terms from the KEGG 2021, Reactome 2022, and Tabula Sapiens databases, based on the top 100 most predictable genes identified by the STimage model. **d**, PCC between STimage-predicted gene expression and ground truth measurements from the Visium experiment in a LOOCV setting, shown for each test sample. **e, f, g**, Tissue plots showing ground truth gene expression for three liver function-related genes from the top 100 most predictable genes. Each row represents a leave-one-out test sample.

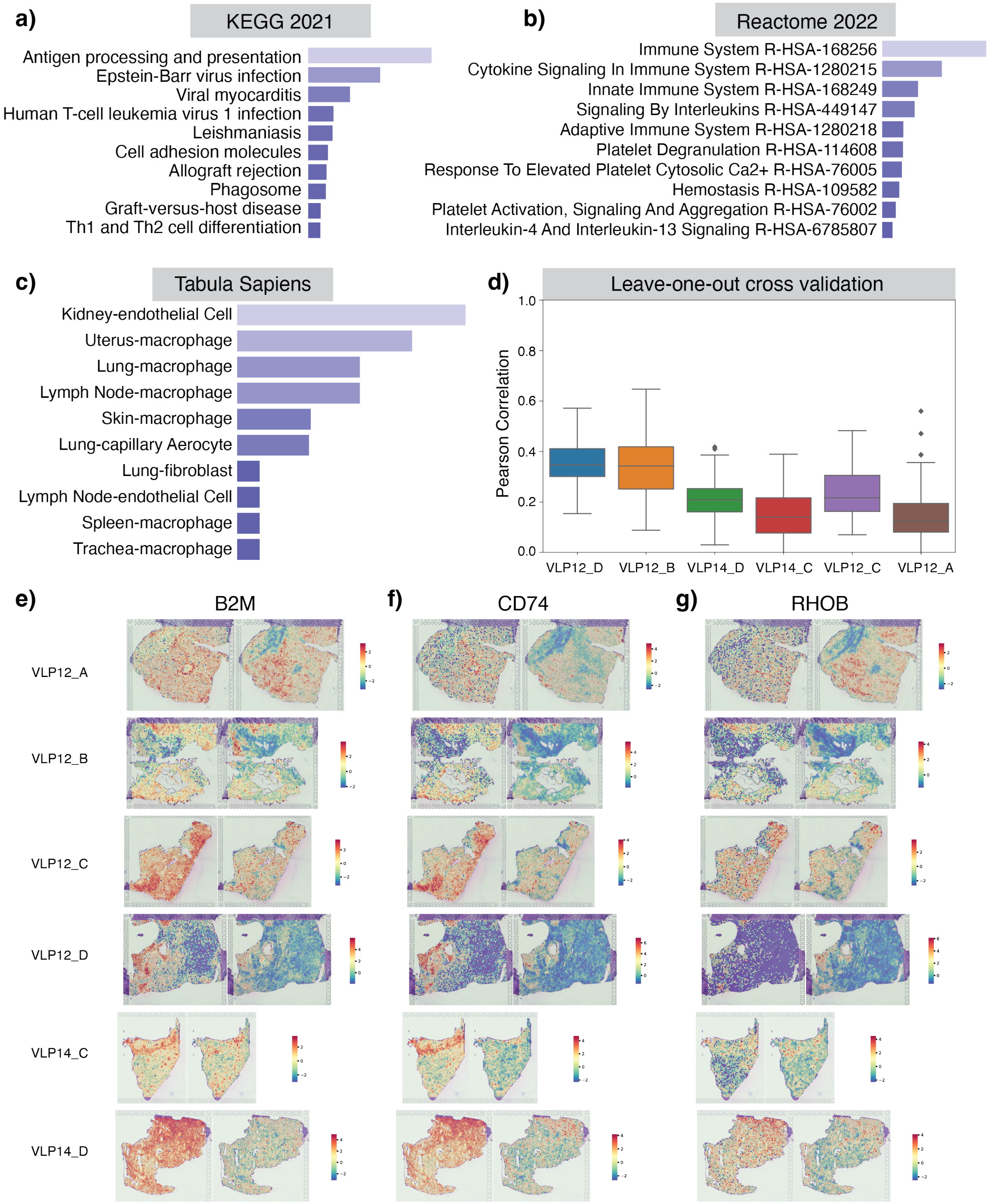


**Figure S7.** Performance of the STimage regression model on a kidney dataset in a LOOCV experiment. **a, b, c**, Top GO enrichment terms from the KEGG 2021, Reactome 2022, and Tabula Sapiens databases, based on the top 100 most predictable genes identified by the STimage model. **d**, PCC between STimage-predicted gene expression and ground truth measurements from the Visium experiment in a LOOCV setting, shown for each test sample. **e, f, g**, Tissue plots showing ground truth gene expression for three kidney function-related genes from the top 100 most predictable genes. Each row represents a leave-one-out test sample.

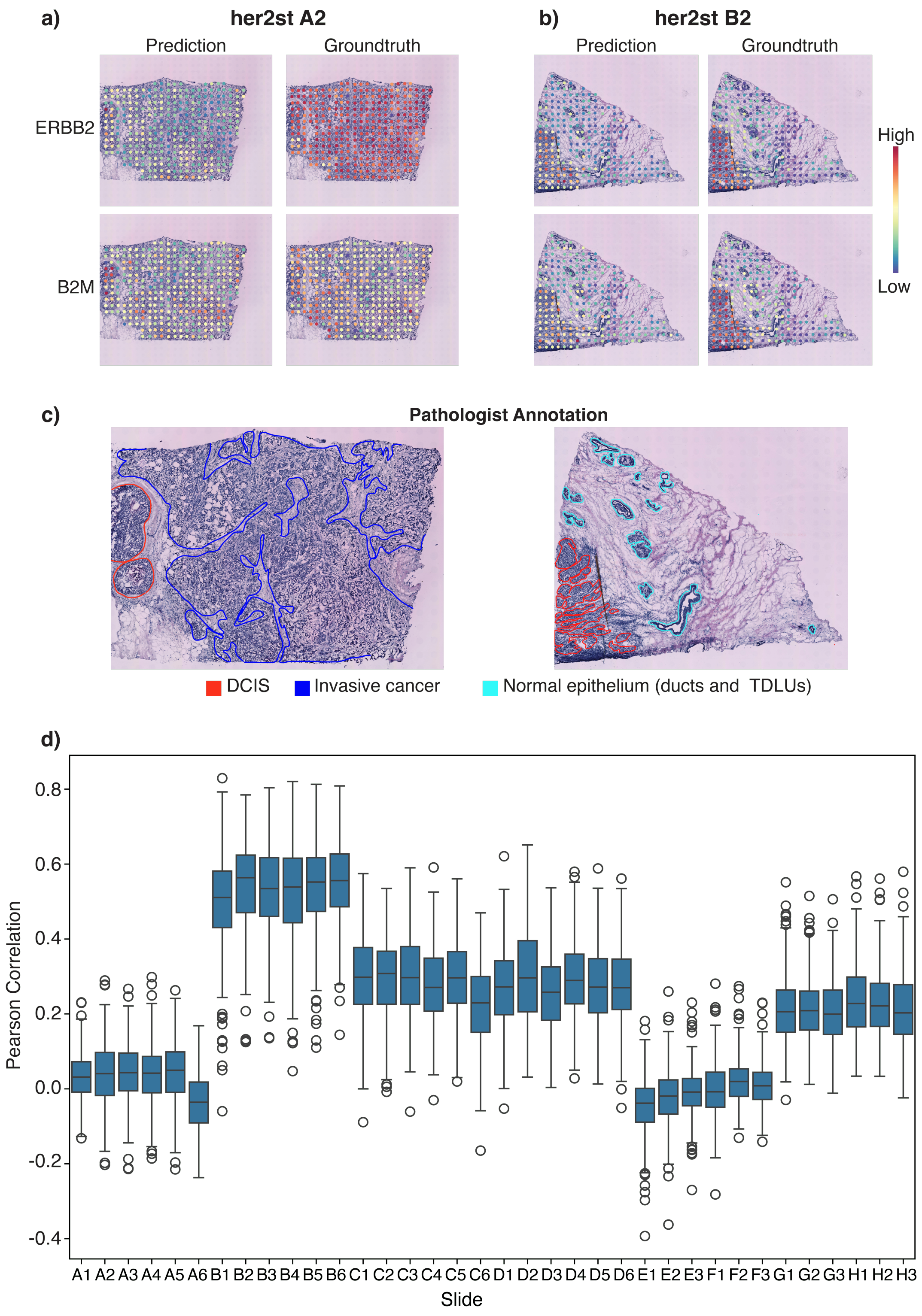


**Figure S8.** Performance evaluation of the model in OOD scenario 1: The model was trained on nine Visium breast cancer datasets and used to predict on the HER2ST legacy ST dataset. **a**, Low-performance sample A2 for two breast cancer marker genes, ERBB2 and B2M. **b**, High-performance sample B2 for the same genes, ERBB2 and B2M. **c**, Pathologist annotations for samples A2 and B2. **d**, Box plots showing the Pearson correlation coefficient (PCC) of 1,146 predicted genes (based on the intersection of the 1,522 predictable genes with those measured in the legacy ST data) compared to ground truth measurements.

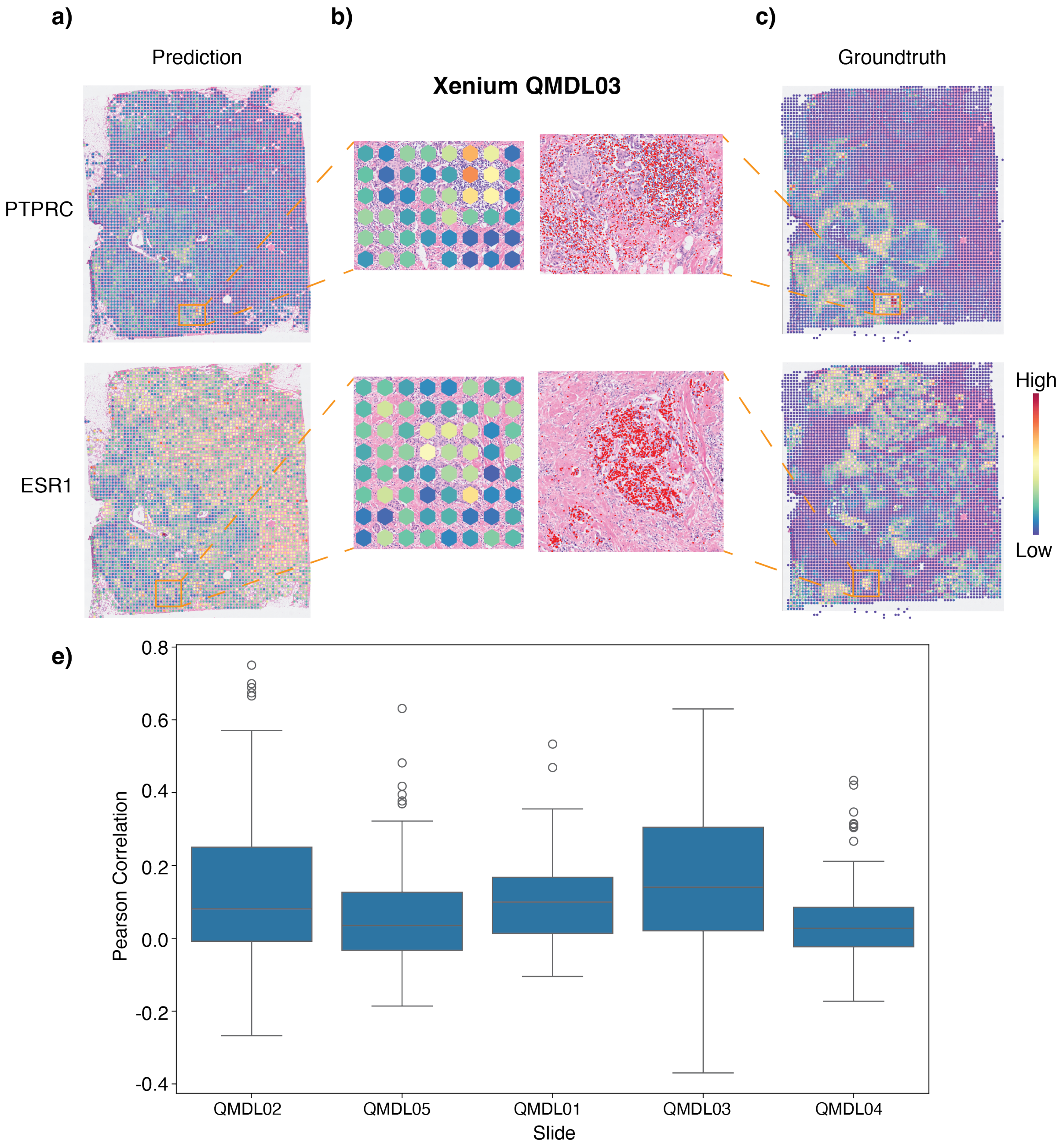


**Figure S9.** Performance evaluation of the model in OOD scenario 2: The model was trained on nine Visium breast cancer datasets and used to predict gene expression in the 10x Xenium breast cancer dataset. **a–c**, Model performance for two breast cancer marker genes, PTPRC and ESR1, in sample QMDL03. **a, c**, Spatial plots of predicted gene expression and ground truth measurements, binned to 55 µm Visium spot-equivalent resolution. **b**, Zoomed-in regions showing detailed visualisation of predictions at the spot level and Xenium gene expression at subcellular resolution. **e**, Box plots showing the Pearson correlation coefficient (PCC) for 106 predicted genes (based on the intersection of 1,522 predictable genes with those measured in the Xenium dataset) compared to ground truth measurements.

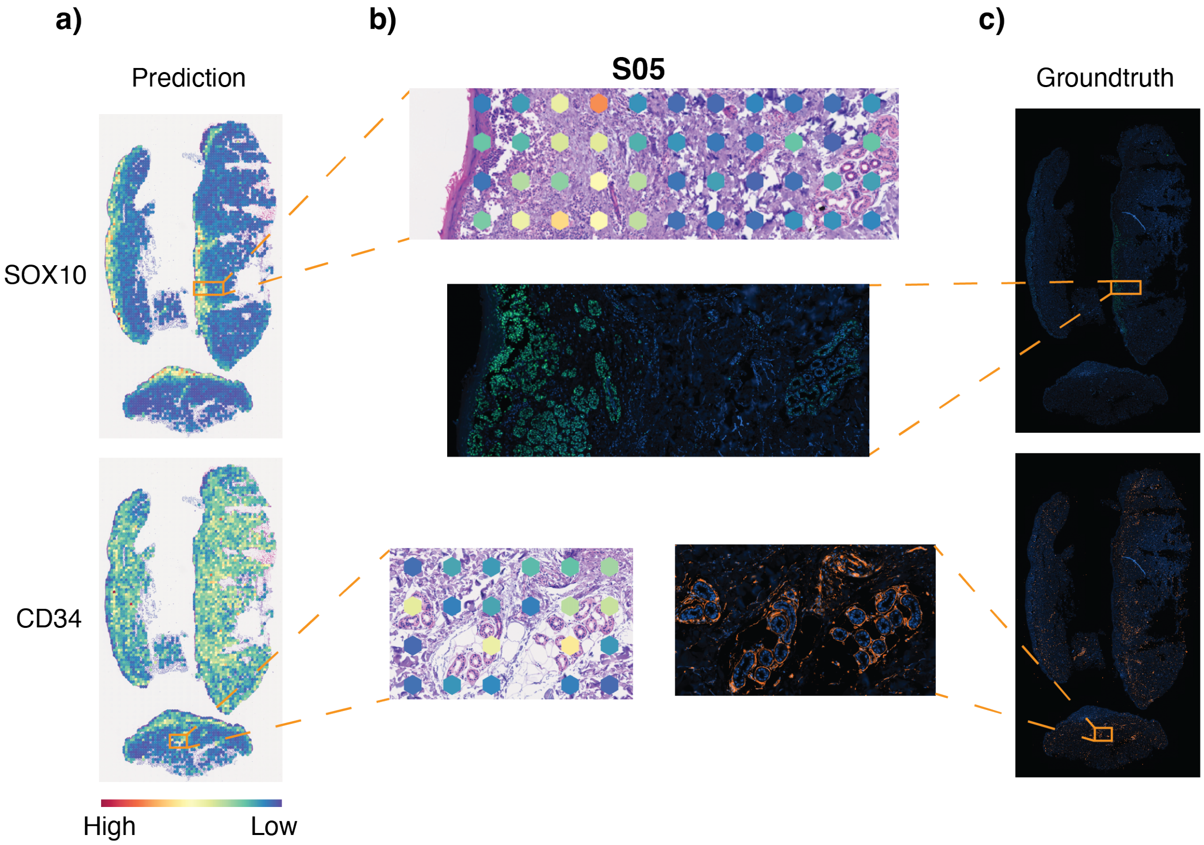


**Figure S10.** Performance evaluation of the model in OOD scenario 3: The model was trained on nine Visium breast cancer datasets and used to predict gene expression in the CODEX spatial proteomics dataset of skin cancer. **a–c**, Spatial plots of predicted gene expression overlaid on the H&E image of the CODEX data (cropped to 55 µm Visium spot-equivalent resolution), alongside ground truth measurements of the CODEX protein markers SOX10 and CD34. **b**, Zoomed-in regions showing detailed visualisation of the model's predictions at the spot level and corresponding CODEX protein expression at subcellular resolution.

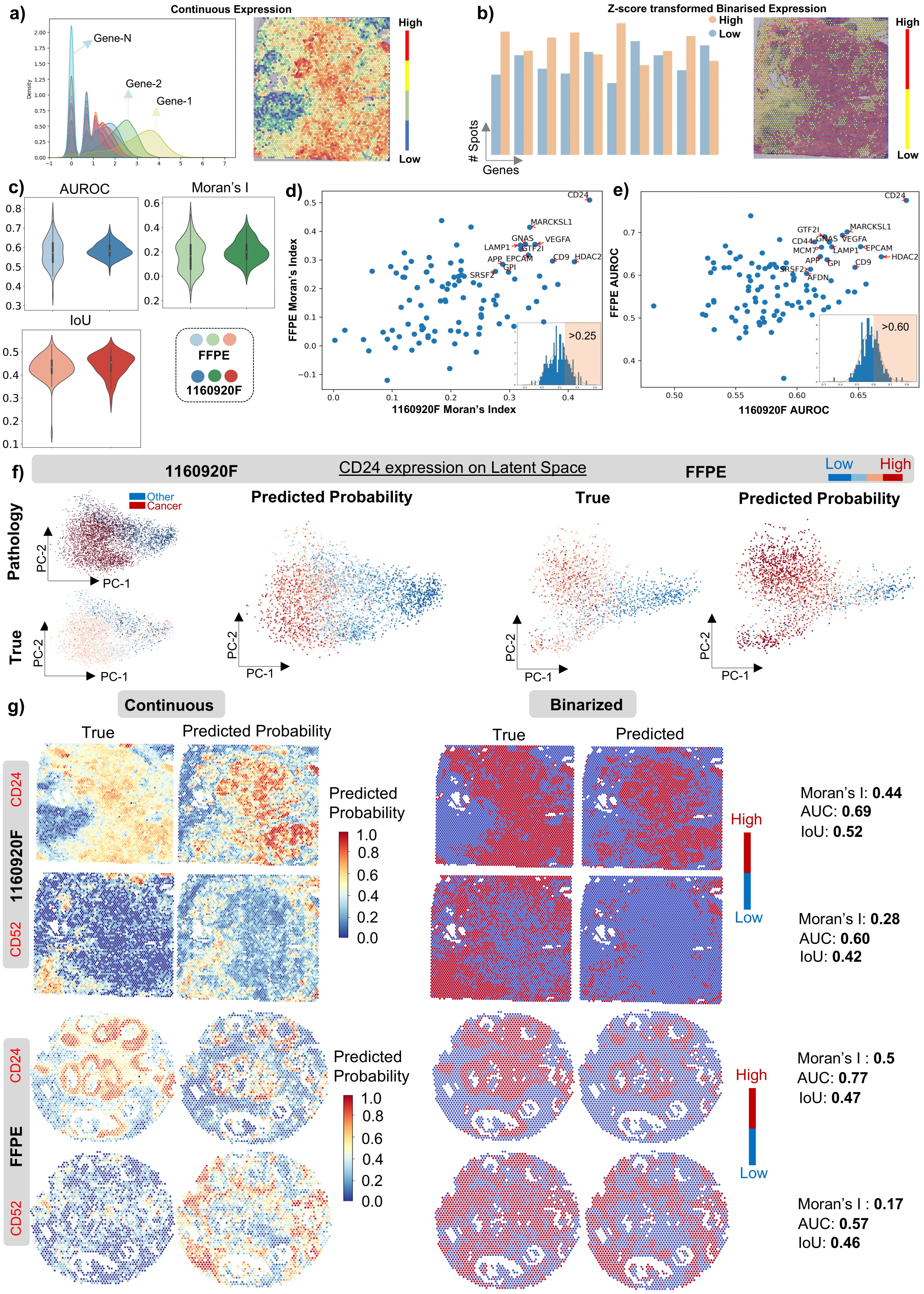


**Figure S11.** The STimage classification model classifies each spot into either a 'High' or 'Low' expression class based on the expression of a given gene. **a**, Density plots of raw gene expression for 10 randomly selected genes, along with an example of spatial gene expression. **b**, Bar plot of z-scored binarised (High/Low) gene expression and an example of binarised spatial gene expression. **c**, AUROC, Moran's I, and IoU scores for the top 100 predictable genes (derived from the regression model) across two test datasets. **d**, Genes with Moran's I > 0.25, and **e**, Genes with AUROC > 0.6 in both datasets, are annotated on the scatter plot (x-axis: 1160920F; y-axis: FFPE). **f**, Visualisation of true gene expression and predicted class probability (High) for the most predictable gene, VEGFA, projected onto the PCA-reduced latent space of ResNet50 features. Each dot represents a Visium spot. The predicted probability aligns with true gene expression and pathologist annotation for 1160920F. **g**, Ground truth and predicted spatial gene expression plots for two complementary genes—CD24 (high in cancer regions) and CD52 (high in immune regions).

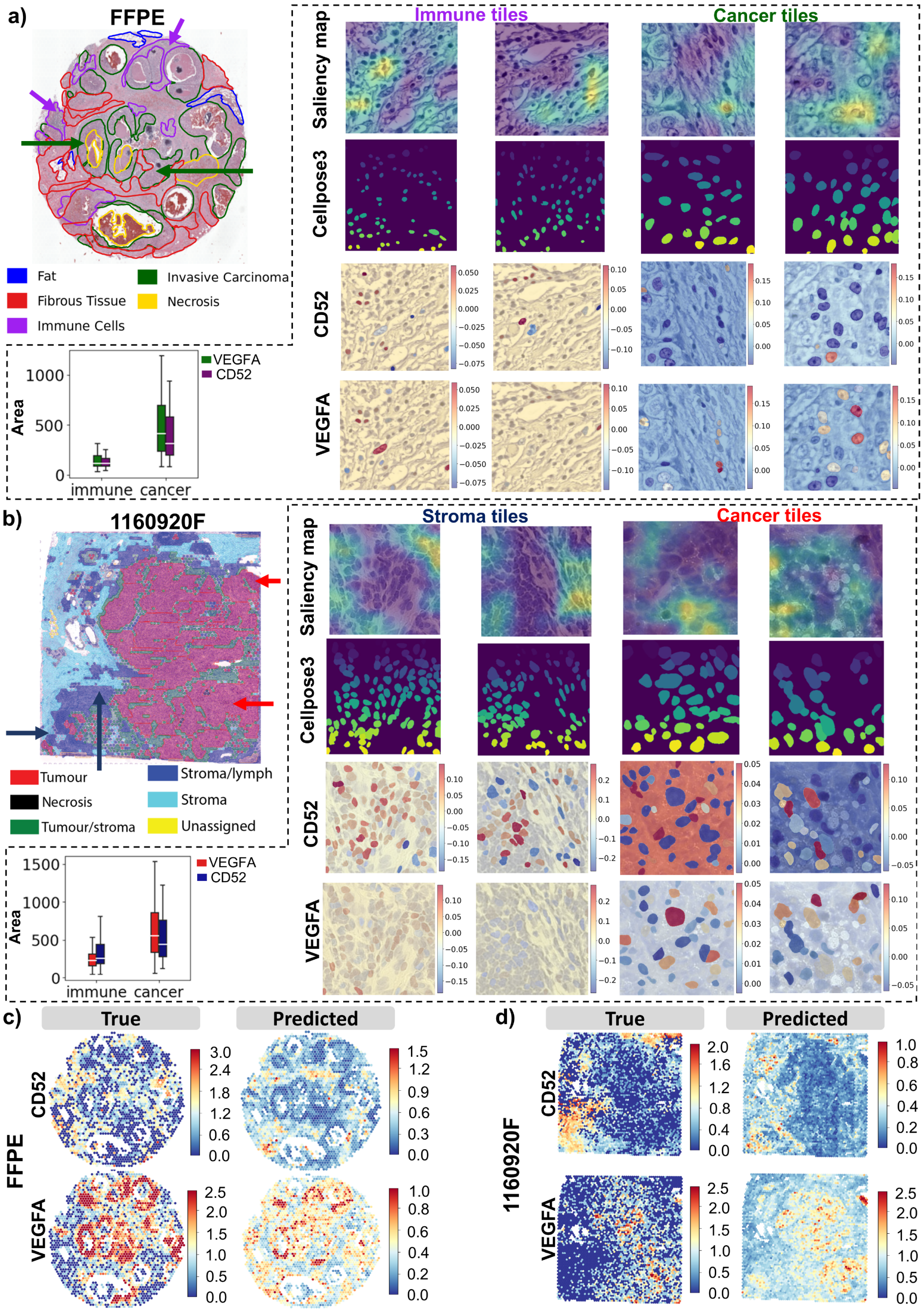


**Figure S12.** STimage interpretability. **a, b**, Immune and carcinoma regions, based on pathologist annotations for two test datasets (FFPE and 1160920F), were selected for interpretability analysis. Arrows indicate the locations of selected tiles in the whole-slide image (WSI), with arrow colours corresponding to the category of the image tile. The first row shows the saliency map from the final convolutional block of ResNet50, demonstrating that ResNet50 learns from the most meaningful features (nuclei) in the image. The second row shows Cellpose-3 segmentation results, which are used as input to the LIME model for interpretability. The third and fourth rows present LIME scores for each nucleus overlaid on the H&E image for immune (CD52) and cancer-related (VEGFA) genes, respectively. Red-coloured nuclei indicate high importance for predicting high expression of the corresponding gene. Immune and stromal tiles in **a, b** show more nuclei with high LIME scores for CD52 compared to VEGFA in both test datasets. Conversely, cancer tiles in **a, b** show more orange-red nuclei with high LIME scores for VEGFA than for CD52. Nuclei contributing most to gene prediction differed between CD52 and VEGFA and between pathological regions. The cancer nuclei (VEGFA, red nuclei) in cancer regions also exhibited larger areas than those with high LIME scores for CD52. **c, d**, Ground truth and STimage-predicted spatial gene expression plots for FFPE and 1160920F, respectively.

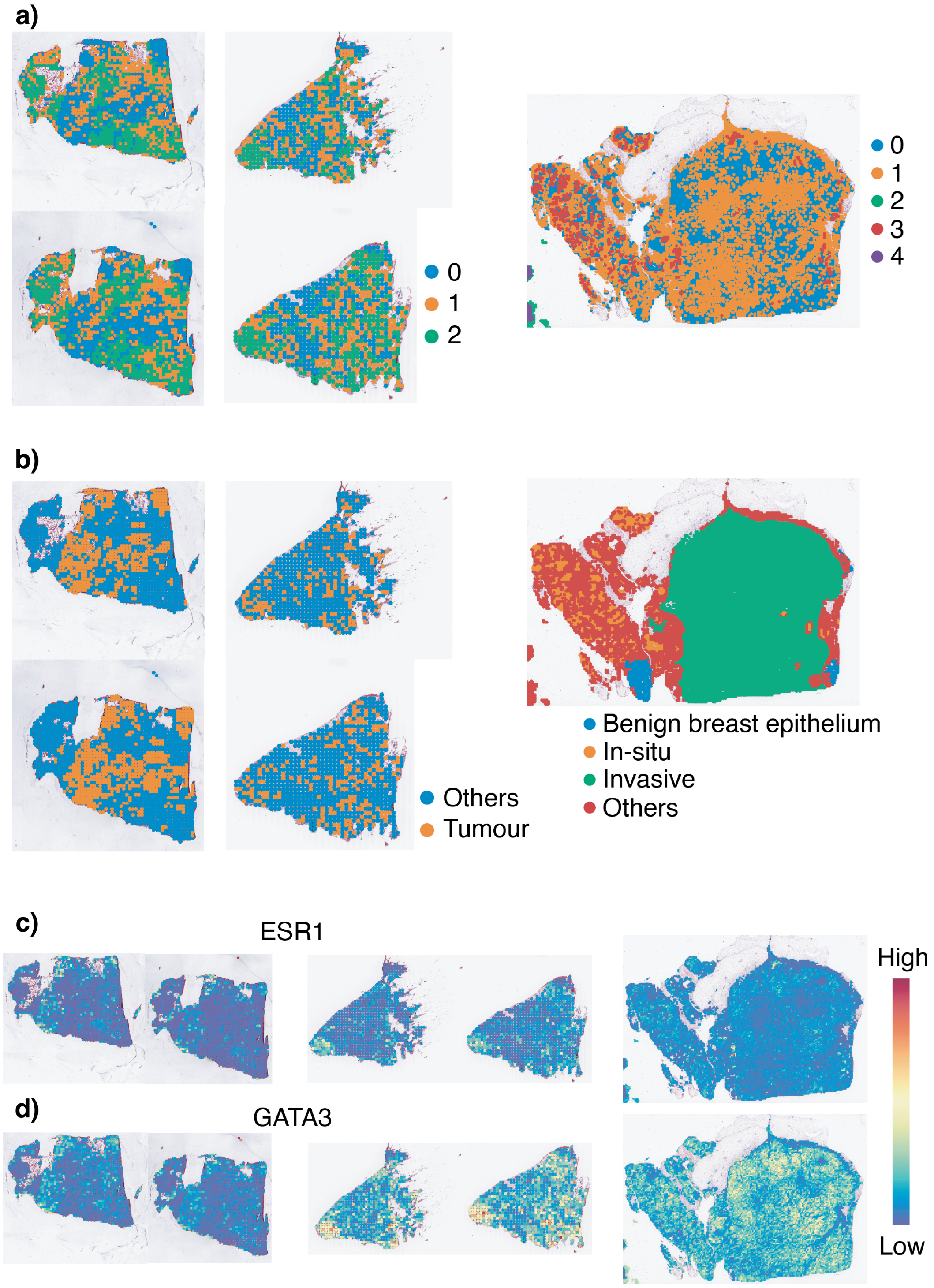


**Figure S13.** Model prediction on external TCGA dataset. This figure presents model predictions for three TCGA breast cancer samples using a model trained on nine Visium breast cancer datasets for 1,522 functional genes. **a**, Louvain clustering based on predicted expression of 1,522 genes for each tile, cropped using a sliding window approach. Each column represents one TCGA sample; the first and second samples include two replicates. **b**, Pathological annotations for each sample. For the first sample (left), Clusters 1 and 2 correspond to tumour regions. For the second sample (middle), Cluster 2 matches tumour regions. For the third sample (right), Cluster 3 corresponds to in situ carcinoma. **c, d**, Spatial plots of two cancer markers, ESR1 and GATA3, from the top-ranked genes, showing strong correlation with pathologist annotations.

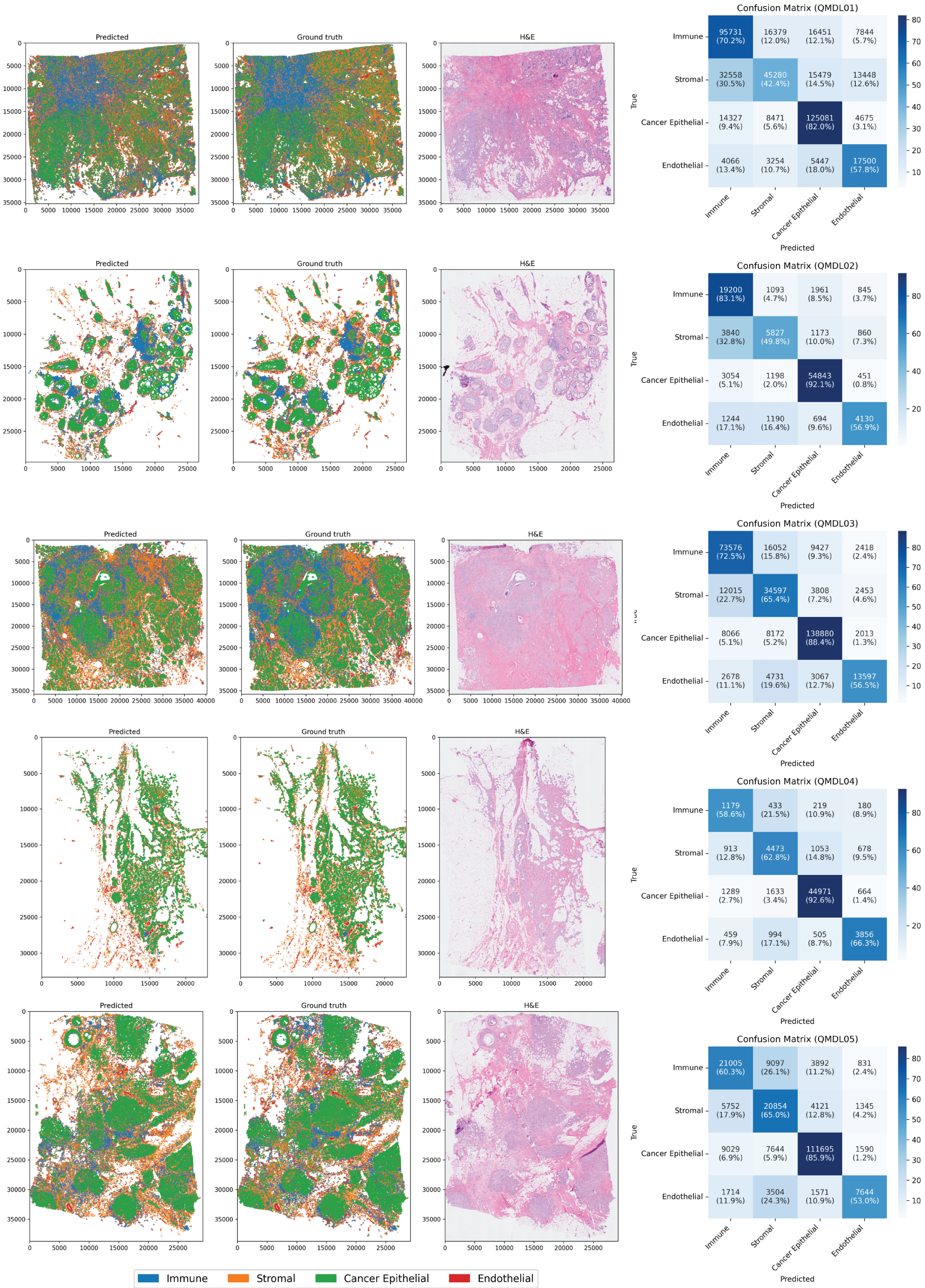


**Figure S14.** Leave-one-out cross-validation for single-cell segmentation and cell type classification in the breast cancer Xenium dataset. Each row represents one test sample in each cross-validation experiment. The first column shows the predicted cell type, the next column contains the ground truth cell type labels from the Xenium data, and the next row presents the HE image, which is the only data required in the prediction step. The last column shows the confusion matrix for all five cell types. The numbers indicate the number of cells, and the percentages represent the proportion of each prediction relative to all ground truth values.

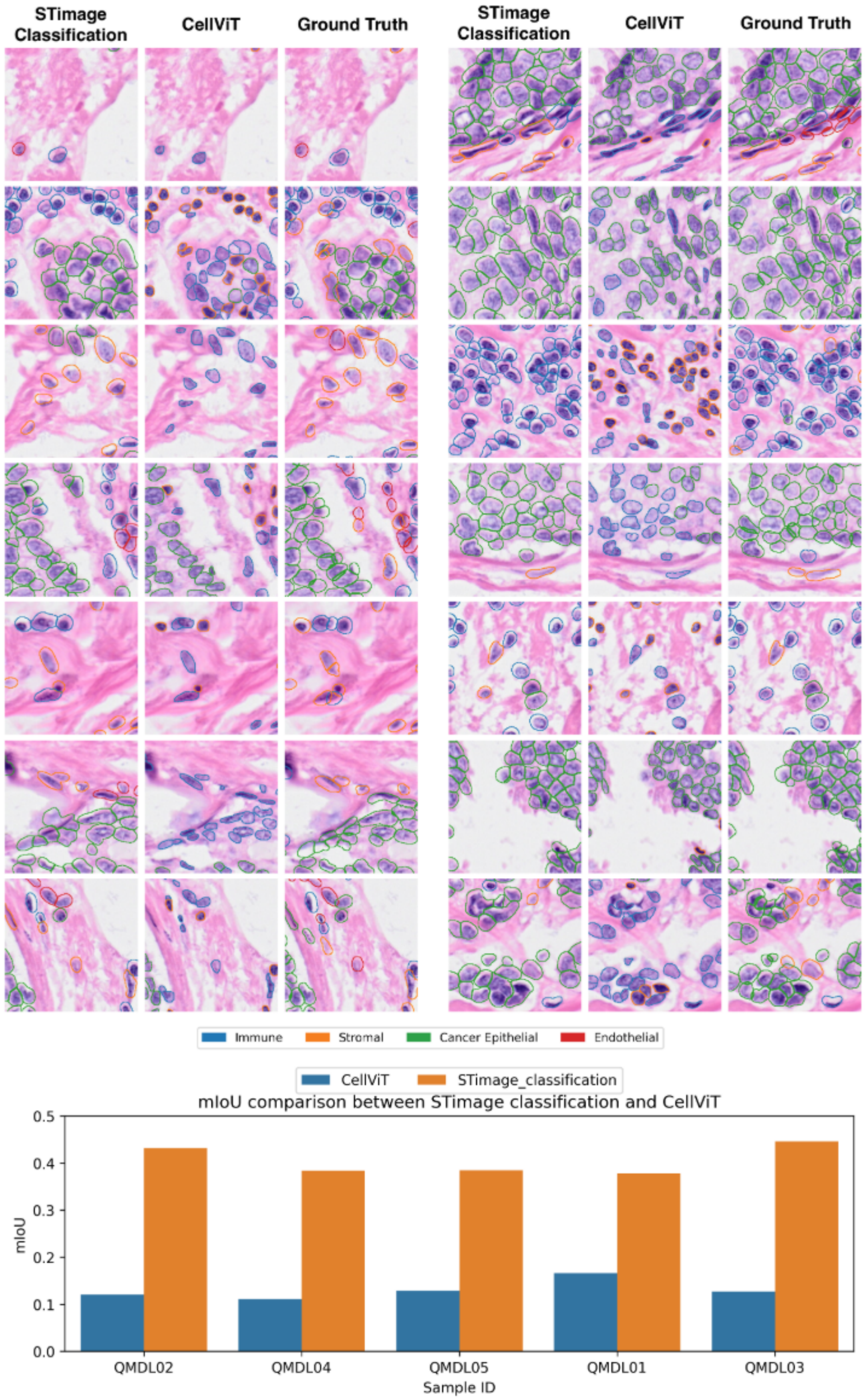


**Figure S15.** The comparison of STimage classification against CellViT in a LOOCV on breast cancer Xenium data. **a**, Randomly selected predictions at the tile level to compare the performance of the STimage classification model, CellViT. Each tile has three columns: from left to right, the STimage classification model, CellViT, and the ground truth. **b**, Mean IoU score for each sample in the LOOCV experiment. The blue bar represents CellViT, while the orange bar represents STimage Classification.

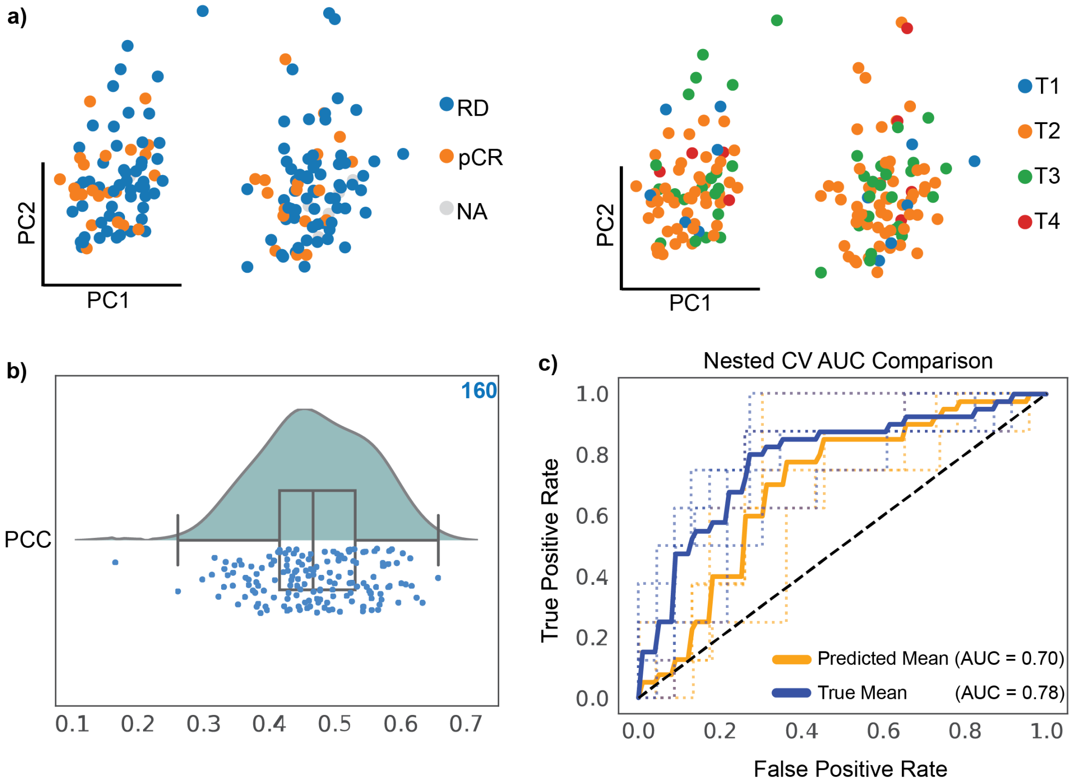


**Figure S16.** Assessing the ability of STimage prediction to stratify patient responses to drug treatment. **a**, The patient cohort with response status on the left [complete response (pCR), residual disease (RD) and not applicable (NA)] and tumour stage on the right (from stage T1 to T4). **b**, Assessing the regression model using PCC. The box plot represents gene-wise PCC per sample, computed between predicted gene expression from H&E images and matched bulk gene expression across 160 samples. Each dot represents a sample. **c**, AUC comparison of true and predicted gene expression to classify patient response to drugs using SVM classifier.

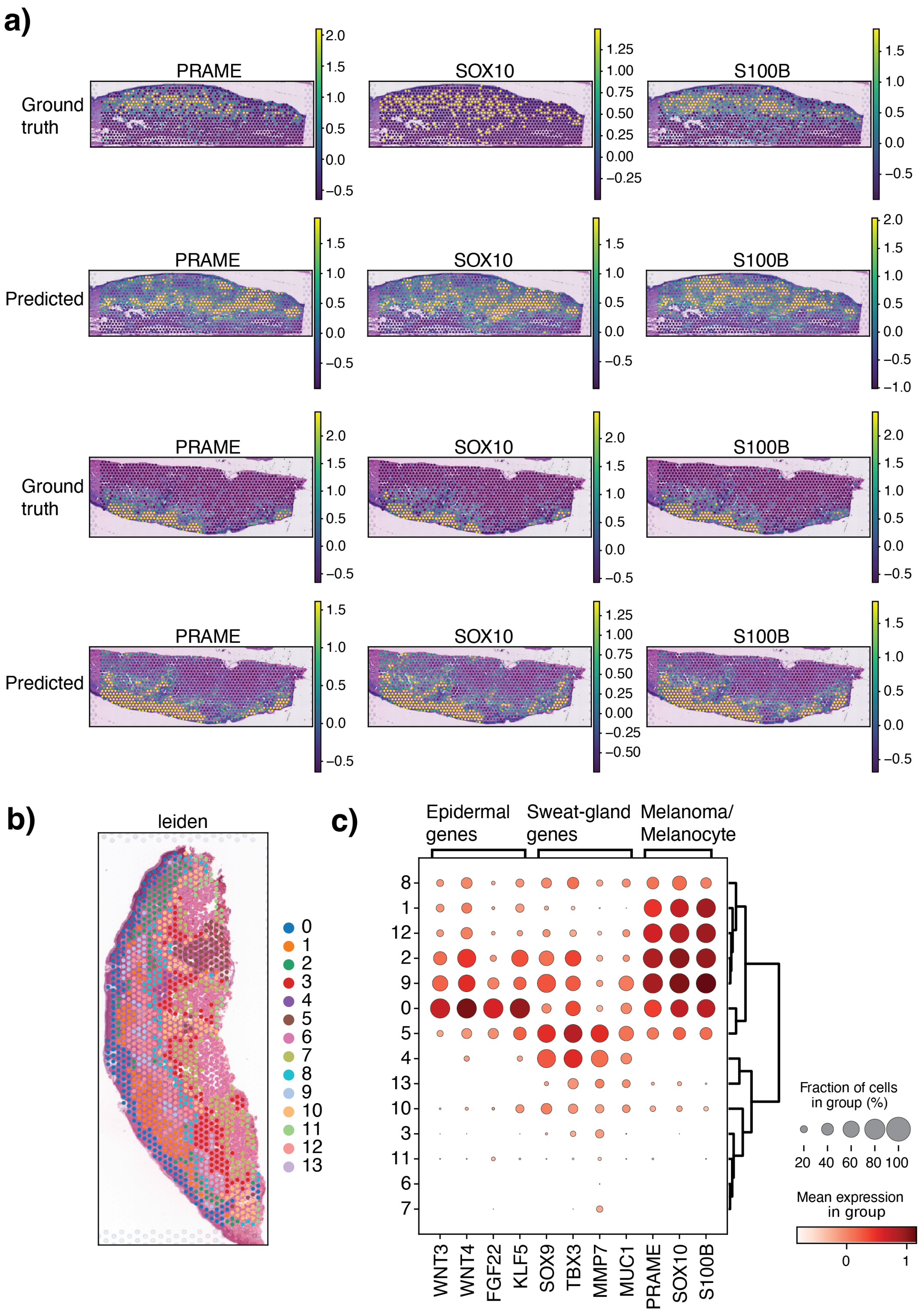


**Figure S17.** Assessing the prediction of biologically relevant markers and tissue regions. **a**, Prediction of the three markers for melanoma, PRAME, SOX10 and S100B. Two samples are shown. For each sample, the ground truth values are from spatial transcriptomics, while the predicted are the results from STimage prediction. **b** and **c** show clustering analysis using predicted values from STimage. Clustering results and gene marker show epidermal layers (cluster 0) and dermal layers with melanocyte/melanoma (1, 2, 9, 12) and sweat gland (clusters 4, 5).
